# Estimating causality between smoking and abdominal obesity by Mendelian randomization

**DOI:** 10.1101/2022.06.06.494971

**Authors:** Germán D. Carrasquilla, Mario García-Ureña, María José Romero Lado, Tuomas O. Kilpeläinen

## Abstract

**Background and Aims:** Smokers tend to have a lower body weight than non-smokers, but also more abdominal fat. It remains unclear whether the relationship between smoking and abdominal obesity is causal. Previous Mendelian randomization studies have investigated this relationship by relying on a single genetic variant for smoking heaviness. This approach is sensitive to pleiotropic effects and may produce imprecise causal estimates. We aimed to assess causality between smoking and abdominal obesity using multiple genetic instruments.

**Methods:** We used GWAS results for smoking initiation (n=1,232,091), lifetime smoking (n=462,690) and smoking heaviness (n=337,334) as exposure traits, and waist-hip ratio (WHR) and waist and hip circumferences (WC and HC) (n up to 697,734), with and without adjustment for body mass index (adjBMI), as outcome traits. We implemented Mendelian randomization using the CAUSE and LHC-MR methods that instrument smoking using genome-wide data.

**Results:** Both CAUSE and LHC-MR indicated a positive causal effect of smoking initiation on WHR (0.13 [95%CI 0.10, 0.16] and 0.49 [0.41, 0.57], respectively) and WHR_adjBMI_ (0.07 [0.03, 0.10] and 0.31 [0.26, 0.37]). Similarly, they indicated a positive causal effect of lifetime smoking on WHR (0.35 [0.29, 0.41] and 0.44 [0.38, 0.51]) and WHR_adjBMI_ (0.18 [0.13, 0.24] and 0.26 [0.20, 0.31]). In follow-up analyses, smoking particularly increased visceral fat. There was no evidence of a mediating role by cortisol or sex hormones.

**Conclusions:** Smoking initiation and higher lifetime smoking may lead to abdominal fat distribution. The increase in abdominal fat due to smoking was characterized by an increase in visceral fat. Thus, efforts to prevent and cease smoking can have the added benefit of reducing abdominal fat.

## INTRODUCTION

Smoking prevention and cessation are critical for public health efforts to reduce the incidence of several chronic disorders, particularly respiratory and cardiovascular diseases [1]. However, smoking cessation is often associated with weight gain [2, 3], which can decrease motivation for sustained cessation and undermine the health benefits. Despite having lower body weight, smokers often have more abdominal fat than non-smokers, which may increase their risk of cardiometabolic diseases [2, 4-13]. Whether the association between smoking and body fat distribution is causal or explained by confounding or reverse causality is still unclear.

Mendelian randomization uses genetic variants associated with exposure traits as instrumental variables to assess whether their relationship with outcome traits may be causal. As maternal and paternal alleles are randomly allocated during conception, Mendelian randomization is analogous to a naturally randomized controlled trial **(Figure 1)**. Three previous Mendelian randomization studies have examined the causal effect of smoking heaviness on abdominal obesity using a single genetic variant in the *CHRNA3/5* smoking heaviness locus [14-16]. The first two studies found no evidence of causality, while the third and largest study suggested a causal relationship between a higher number of cigarettes per day and higher WHR, even after adjusting for BMI [15].

**Figure 1:**
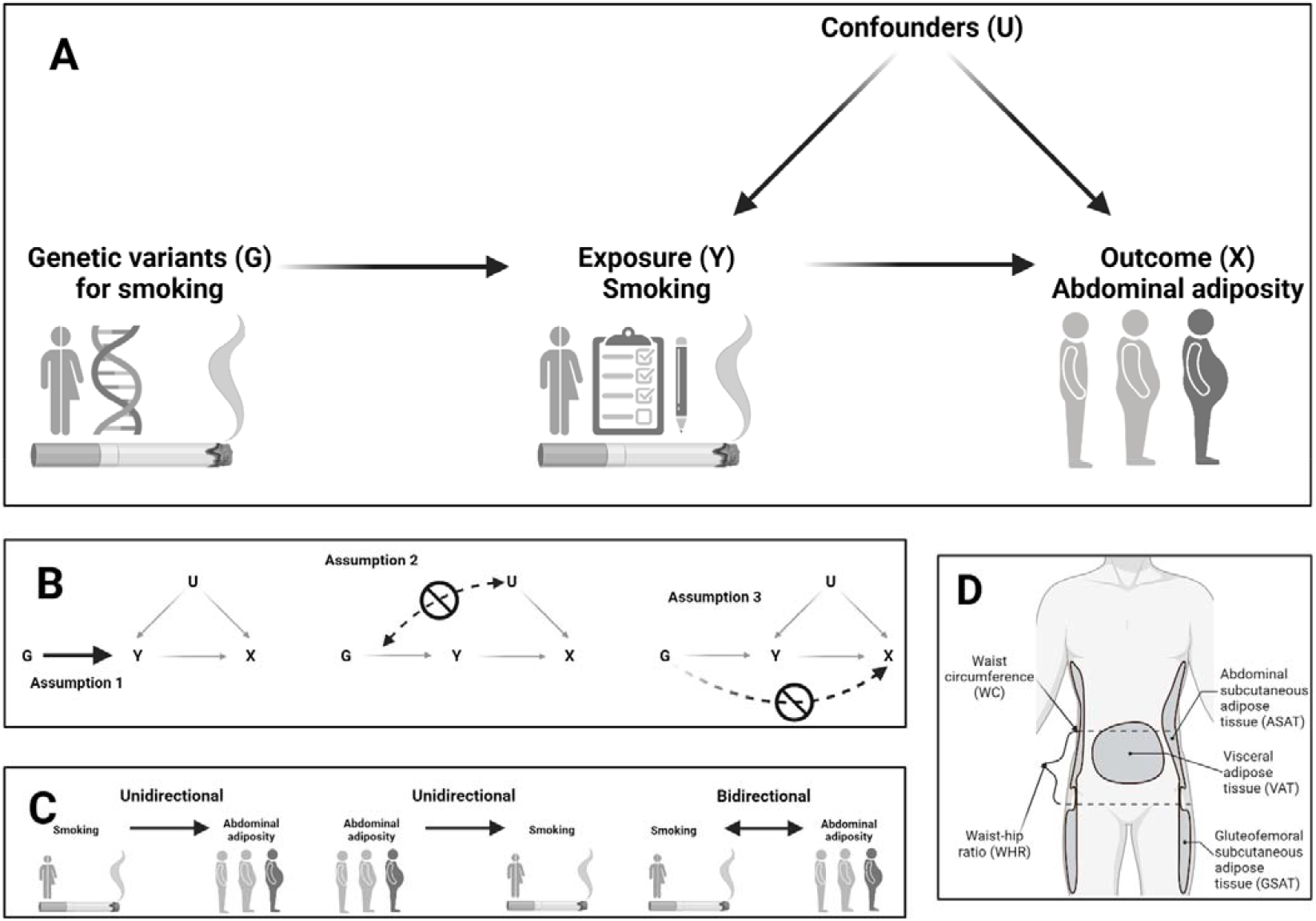
Methodological considerations for examining causality between smoking and abdominal adiposity. Mendelian randomization can provide an unbiased estimation of causality (A), provided that three assumptions are valid (B). The first assumption is that the instrument is associated with the exposure. The second assumption is that the instrument is not associated with the outcome through a confounding pathway (uncorrelated horizontal pleiotropy). The third assumption is that the instrument does not directly influence the outcome but does so indirectly through the exposure (correlated horizontal pleiotropy). Depending on the direction of the causal estimate being tested, the analysis can be unidirectional or bidirectional (C). Abdominal adipose tissue consists of visceral adipose tissue and abdominal subcutaneous adipose tissue (D).

Studies based on a single genetic instrument are sensitive to genetic pleiotropy and may produce imprecise causal estimates, leaving the relationship between smoking and abdominal obesity uncertain. Since the publication of the previous Mendelian randomization studies, genome-wide association studies (GWAS) of smoking traits have identified >400 novel loci associated with smoking traits [17]. These findings provide new opportunities for increasing the precision and statistical power of Mendelian randomization studies while reducing bias. Furthermore, new and improved Mendelian randomization methods allow leveraging GWAS summary results at genome-wide level to better control for pleiotropy and latent heritable confounders and assess bidirectional causal effects [18-20].

Here, we report on two-sample Mendelian randomization analyses to estimate the causal effect of smoking initiation, smoking heaviness, and lifetime smoking (which captures smoking heaviness, duration, and time of cessation) on abdominal adiposity.

## METHODS

### Mendelian randomization methods

The Causal Analysis Using the Summary Effect estimates (CAUSE) and the Latent Heritable Confounder MR (LHC-MR) methods utilize genome-wide association data to assess causal relationships rather than genome-wide significant loci only, to correct for sample overlap and better control for correlated and uncorrelated horizontal pleiotropy (**Figure 1, B**) [19, 20]. The CAUSE method assesses the unidirectional causal relationship (**Figure 1, C**) between an exposure and an outcome trait by calculating the posterior probabilities of a causal effect and a shared effect. The causal effect reflects the effect of the variants on the outcome trait through the exposure, while the shared effect reflects the effect of the variants on the outcome trait through confounders (correlated horizontal pleiotropy, **Figure 1, B**). The LHC-MR method assesses bidirectional causal relationships (**Figure 1, C**) by dividing the association between an exposure and an outcome trait into four different effects: the causal effect of the exposure on the outcome, the causal effect of the outcome on the exposure, the effect of confounders that affect the outcome through the exposure (vertical pleiotropy), and the effect of confounders that affect the outcome independently of the exposure (correlated horizontal pleiotropy) [20]. More details on the CAUSE and LHC-MR methods are provided in the **Supplementary Note**.

In addition to the CAUSE (v1.0.0) and LHC-MR (v.1.0.0) methods, we implemented Mendelian randomization using inverse variance weighted, MR-Egger, weighted median, and weighted mode methods, instrumenting the smoking exposure trait using genome-wide significant loci (P < 5×10^−8^). We selected lead variants that showed a pairwise linkage disequilibrium (LD) r^2^ < 0.001 and a distance of 10.000kb with all other lead variants associated with the same trait. Variants not available in the outcome trait GWAS were substituted by their LD proxies (r^2^ > 0.8). Ambiguous palindromic variants (A/T, G/C) were excluded. We applied Steiger filtering to remove variants that are likely to affect the outcome trait through other traits than the exposure trait. The strength of the genetic instruments was assessed using the F statistic. The analyses were performed using the TwoSampleMR (v.0.5.6) package in R [18].

We estimated heterogeneity across the causal estimates of SNPs using the Meta R package [23]. The causal estimates were considered heterogeneous if the P value for Cochran’s Q test was significant (P<0.05) and I^2^ was above 25%. We assessed bias introduced by horizontal pleiotropy by implementing the Egger’s intercept test. Egger’s intercept P<0.05 was considered as evidence of horizontal pleiotropy. We used the Rucker framework [24] to assess whether MR-Egger regression, which accounts for horizontal pleiotropy but limits statistical power, should be applied instead of the standard inverse variance-weighted model. We visually assessed heterogeneity and horizontal pleiotropy using leave-one-out forest plots and funnel plots. To detect individual pleiotropic variants that might bias the results, we used the RadialMR package in R (1.0.0) [25], applying an iterative Cochran’s Q method and setting a strict outlier threshold of P<0.05. The iterative Cochran’s Q, either inverse variance-weighted Q or Egger Q, was chosen depending on the Rucker framework results. After removing outlier variants detected with RadialMR, we re-ran the Mendelian randomization and sensitivity tests to ensure that horizontal pleiotropy introduced by the outlier variants had been removed.

There is a considerable genetic correlation between smoking status and socioeconomic status [26]. Therefore, we conducted a multivariable Mendelian randomization (MVMR, v. 0.3) [27] analysis to assess the causal effects of smoking initiation and lifetime smoking on abdominal adiposity while controlling for the effect of socioeconomic status (educational attainment, n=458,079) [28]. We followed the MVMR steps described previously by Sanderson et al [29]. First, we identified independent, genome-wide significant variants associated with smoking and education (r^2^ < 0.001 within 10.000kb distance) and replaced them with proxies (r^2^ > 0.8) when not available in the outcome trait GWAS. We calculated the conditional mean F-statistic to ensure that the instrumental variable was robustly associated with the exposure trait. We obtained IVW estimates before and after outlier removal for genetic instruments with a mean F statistic > 10. We assessed pleiotropy with conditional Cochran’s Q, where P<0.05 was interpreted as evidence of pleiotropy. Further details on the MVMR procedures can be found in the **Supplementary Note**.

This study has been reported according to the STROBE-MR guidelines [30] found at the end of the **Supplementary Note**.

### Data sources for smoking and body fat distribution

We utilized the largest published GWAS summary level data of European ancestry for smoking initiation (n=1,232,091), defined as whether an individual has ever smoked regularly (yes/no) [17]; lifetime smoking (n=462,690), which captures the initiation, duration, heaviness, and time since the cessation of smoking [31]; and smoking heaviness, defined as cigarettes smoked per day by current smokers only or current and former smokers combined (n=337,334) [17], or as the total number of pack-years smoked in adulthood (n = 142,387) (**Supplementary Table 1**).

We also utilized the largest published GWAS summary level data of European ancestry for measures of body fat distribution (**Figure 1, D**), including WHR (n=697,734) and WHR adjusted for body mass index (WHR_adjBMI_, n=694,649) from a meta-analysis of the Genetic Investigation of Anthropometric Traits (GIANT) consortium and the UK Biobank [32]; waist and hip circumferences (WC and HC) from the UK Biobank (n=462,166 and n=462,177, respectively); and WC_adjBMI_ and HC_adjBMI_ from the GIANT consortium (n=231,355 and n=211,117, respectively) [33]. In the analyses of the causal relation between cigarettes smoked per day and abdominal adiposity in current smokers, we utilized the GIANT consortium results for WHR_AdjBMI_ and WC_AdjBMI_ in current smokers only (n=40,543 and n=43,226, respectively) [33]. In the Mendelian randomization analyses using the IVW, MR Egger, weighted median and weighted mode methods, which are sensitive to bias when there is a sample overlap between the exposure and outcome traits, we used the GIANT consortium data without the UK Biobank for each of the body fat distribution traits (n between 40,543 and 232,101) [34] (**Supplementary Table 1**).

### Association of smoking variants with body fat depot volumes and hormonal levels

To understand the effect of smoking variants on visceral fat (VAT), abdominal subcutaneous fat (ASAT), gluteofemoral subcutaneous fat (GSAT), and their respective ratios VAT/ASAT, VAT/GSAT and ASAT/GSAT (n=38,965) [35] (**Figure 1, D**), we constructed genetic risk scores using variants from the inverse variance-weighted model for the causal association between smoking initiation and WHR (121 variants), smoking initiation and WHR_adjBMI_ (130 variants), lifetime smoking and WHR (76 variants), or lifetime smoking and WHR_adjBMI_ (83 variants). We computed weighted genetic scores using the beta of the variants for the smoking trait as weights with gtx.package’s (v0.0.8) grs.summary function [36]. This function approximates the results of a genetic risk score using GWAS summary statistics by calculating the joint effect of genetic variants on an outcome trait. As previous observational studies suggest that effects of smoking on body fat distribution may be explained by changes in hormones such as cortisol or sex hormones, we also assessed the effect of the genetic risk scores on the levels of cortisol (N=25,314) [37], oestradiol (N=311,675) [38], testosterone (N=425,097) and sex hormone-binding globulin (N=370,125) [39].

The present study used public GWAS summary-level data and did not have direct contact with study participants. Therefore, no direct ethical approval was needed. The code and curated data for the current analysis are available at https://github.com/MarioGuCBMR/MR_smoking_abdominal_adiposity.

## RESULTS

### Effects of smoking initiation and lifetime smoking on abdominal adiposity

In Mendelian randomization analyses by the CAUSE and LHC-MR methods, we found evidence of a positive causal effect of smoking initiation on WHR (0.13 [0.10, 0.16] and 0.49 [0.41, 0.57], respectively) and WHR_adjBMI_ (0.07 [0.03, 0.10] and 0.31 [0.26, 0.37]). Similarly, we found evidence of a positive causal effect of lifetime smoking on WHR (0.35 [0.29, 0.41] and 0.44 [0.38, 0.51]) and WHR_adjBMI_ (0.18 [0.13, 0.24] and 0.26 [0.20, 0.31]) (**Figure 2, Supplementary Tables 2-4, Supplementary Figure 1**). The CAUSE method showed no evidence of correlated horizontal pleiotropy, as indicated by the zero median shared effect and low q-values in all models (**Supplementary Table 2**). The LHC-MR results indicated an effect of confounders on WHR and WHR_adjBMI_, independent of smoking traits, suggesting the presence of correlated horizontal pleiotropy (**Supplementary Tables 4-5**). However, the reported confounder effect was directionally opposite to the positive causal effects of smoking initiation and lifetime smoking on WHR and WHR_adjBMI_, suggesting that the confounder effect leads to an underestimation of the true causal effects.

**Figure 2:**
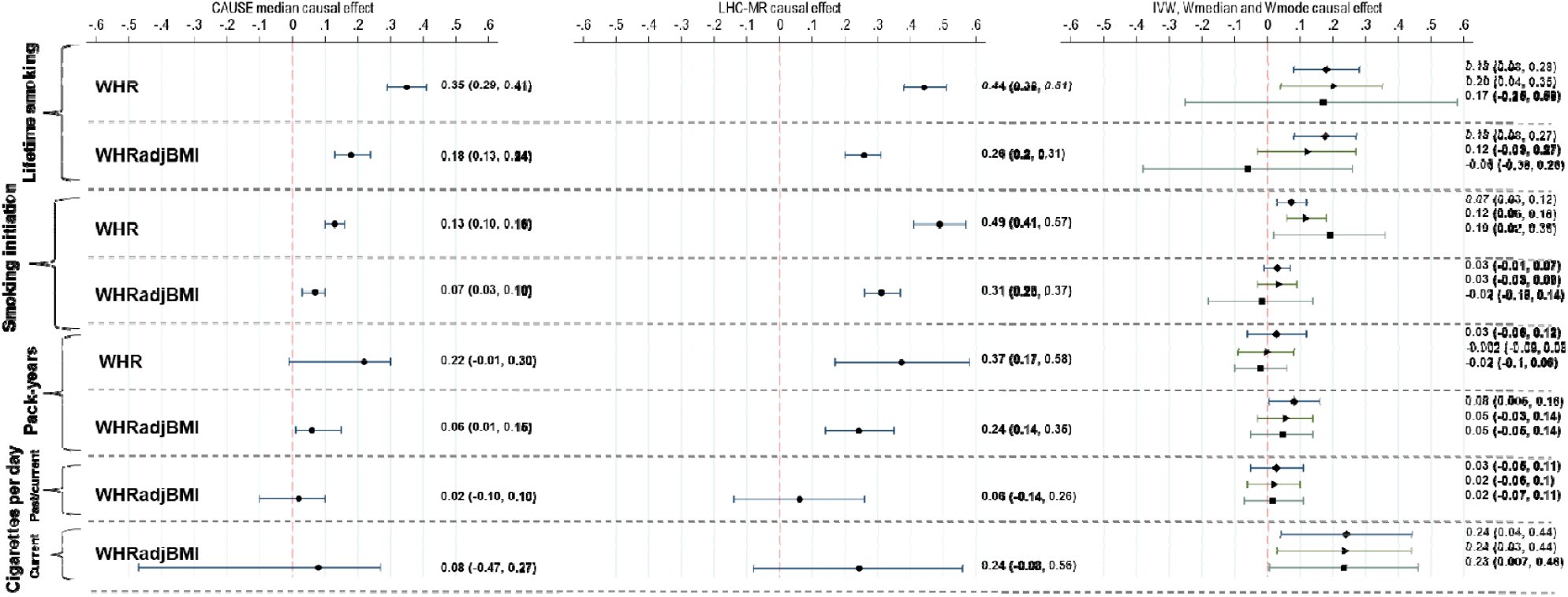
Mendelian randomization results for the causal effect of smoking traits on waist-hip ratio (WHR) and waist-hip ratio adjusted for body mass index (WHR_adjBMI_). The left panel shows the results from the CAUSE method, the middle panel shows the results from the LHC-MR method, and the right panel shows the results from the inverse-variance-weighted (diamond), weighted median (triangle) and weighted mode (square) methods.

As WHR is calculated as the ratio of WC and HC, the causal effect of smoking on WHR could be due to an effect on either WC or HC, reflecting the deposition of abdominal and gluteofemoral fat, respectively. When we examined the causal effects of smoking initiation and lifetime smoking on WC using the CAUSE and LHC-MR methods, we found evidence of a positive causal effect of smoking initiation on WC (0.12 [0.09, 0.16] and 0.37 [0.14, 0.6], respectively) and on WC_adjBMI_ (0.03 [-0.04, 0.11] and 0.07 [0.03, 0.11]), as well as evidence of a positive causal effect of lifetime smoking on WC (0.35 [95%CI 0.29, 0.41] and 0.42 [0.21, 0.63]) and on WC_adjBMI_ (0.09 [-0.05, 0.23] and 0.02 [-0.03, 0.06]) (**Figure 3, Supplementary Tables 2-5**). While there was also evidence of a positive causal effect on HC (**Figure 3, Supplementary Tables 2-5**), the effect estimates were smaller than for WC and changed direction from positive to negative after adjustment for BMI. This suggests that smoking initiation and lifetime smoking may lead to relatively higher abdominal fat storage at the cost of lower gluteofemoral fat.

**Figure 3:**
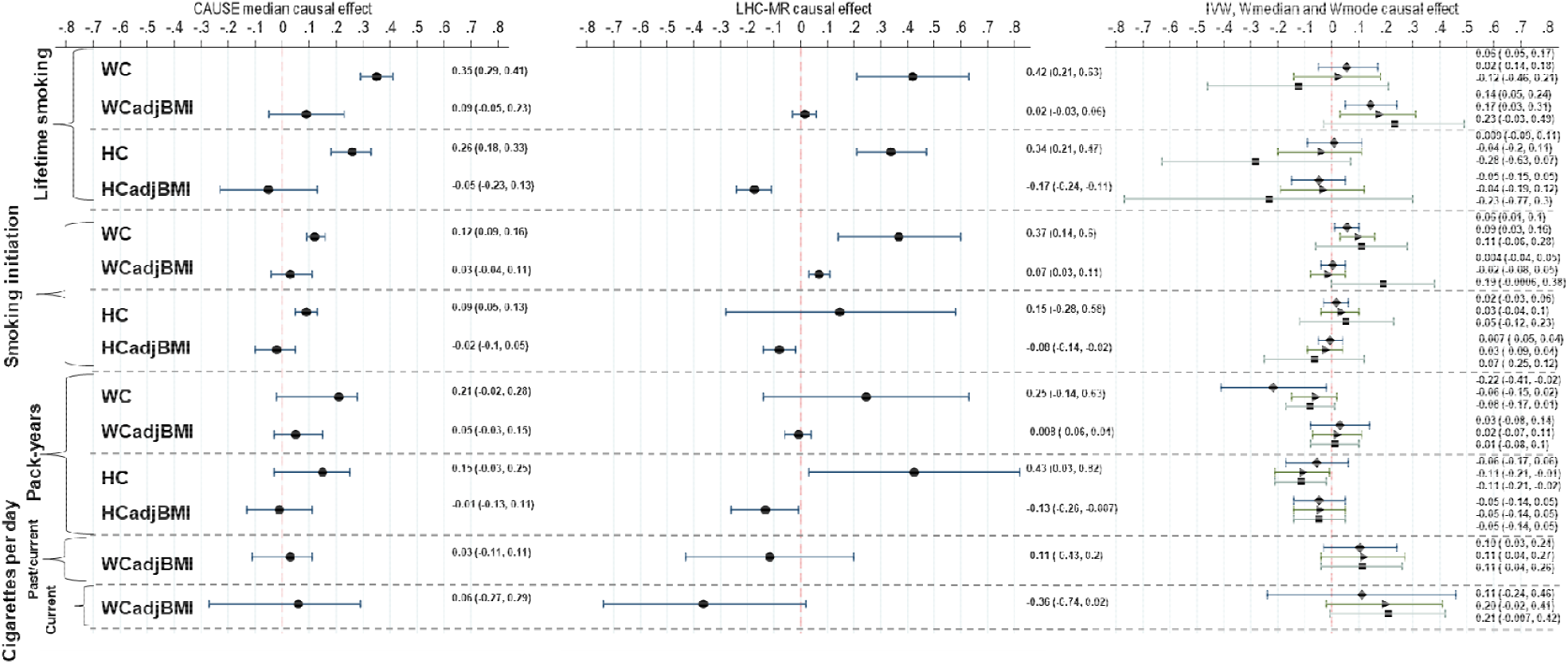
Mendelian randomization results for the causal effect of smoking traits on waist circumference (WC), waist circumference adjusted for body mass index (WC_adjBMI_), hip circumference (HC), and hip circumference adjusted for body mass index (HC_adjBMI_). The left panel shows the results from the CAUSE method, the middle panel shows the results from the LHC-MR method, and the right panel shows the results from the inverse-variance-weighted (diamond), weighted median (triangle) and weighted mode (square) methods.

Using Mendelian randomization methods that instrument the smoking exposure using genome-wide significant variants only (IVW, MR-Egger, weighted median, weighted mode), the results were largely consistent with those from CAUSE and LHC-MR. In particular, there was evidence of causal effects of lifetime smoking and smoking initiation on WHR (IVW: 0.18 [95%CI 0.08, 0.28] and 0.07 [0.03, 0.12], respectively) and WHR_adjBMI_ (IVW: 0.18 [0.08, 0.27] and 0.03 [-0.01, 0.07], respectively) (**Figure 2, Supplementary Tables 6-7, Supplementary Figures 2 and 3**). Similar to CAUSE and LHC-MR, the causal estimates for WC_adjBMI_ and HC_adjBMI_ suggested that higher abdominal fat was at the cost of lower gluteofemoral fat (**Figure 3, Supplementary Tables 6-7, Supplementary Figures 2 and 3**). In original units, our results suggest that starting to smoke increases WHR by an average of 0.063 (95% CI: 0.012-0.114). This corresponds to an increase of approximately 7% in WHR, given that the average WHR in the population is 0.90.

There is a considerable genetic correlation between smoking initiation and markers of socioeconomic status, such as educational attainment (r_g_ ∽-0.4) [26]. To determine whether the causal effects of smoking initiation and lifetime smoking on abdominal obesity are independent of genetic effects on socioeconomic status, we performed a multivariable Mendelian randomization analysis. We found that smoking initiation and lifetime smoking are causally associated with WHR, WHR_adjBMI_, WC and WC_adjBMI_, even after controlling for genetic effects on educational attainment (**Supplementary Note, Supplementary 8-9**). This suggests that the causal effects of smoking initiation and lifetime smoking on abdominal obesity are independent of socioeconomic status.

### Effect of smoking heaviness on abdominal adiposity in past and current smokers

Our Mendelian randomization analysis showed no evidence of a causal relationship between a higher number of cigarettes per day or the cumulative pack years of smoking in adulthood and WHR or WHR_adjBMI_ (**Figure 2-3, Supplementary Tables 2-5**). This suggests that the causal effect of lifetime smoking on abdominal obesity is not related to the heaviness of smoking per se.

A previous Mendelian randomization study found a causal association between smoking heaviness and WHR_adjBMI_ in current smokers when using a single genetic instrument in the *CHRNA3/5* locus [15]. Our results were consistent with this finding when we investigated Wald ratio estimates for *CHRNA3/5* locus alone (**Supplementary Table 10**). The lack of evidence of a causal effect when including all currently known variants for smoking heaviness suggests that the previous findings may have been due to a locus-specific pleiotropic effect on abdominal obesity (**Supplementary Figure 4**).

### Causal effect of abdominal obesity on smoking traits

We used the LHC-MR method to estimate bidirectional causal effects between abdominal obesity and smoking traits. Our results showed a positive causal effect of WHR on lifetime smoking (0.08 [0.04, 0.13]), WC on smoking initiation (0.12 [0.04, 0.21]) and lifetime smoking (0.14 [0.03, 0.25]), and HC on smoking initiation (0.11 [0.03, 0.2]) and lifetime smoking (0.10 [0.04, 0.16]) (**Figure 4, Supplementary Table 5**). However, these effects disappeared after adjusting for BMI, suggesting that they may be due to a causal effect of obesity on smoking, as shown before [40]. For smoking heaviness, we found a causal effect of WHR_adjBMI_ on both cigarettes per day (0.24 [0.11, 0.37]) and the number of smoking pack years in adulthood (0.07 [0.03, 0.12]). This suggests that abdominal adiposity may be causally associated with heavier smoking, independent of BMI (**Figure 4, Supplementary Table 5**).

**Figure 4:**
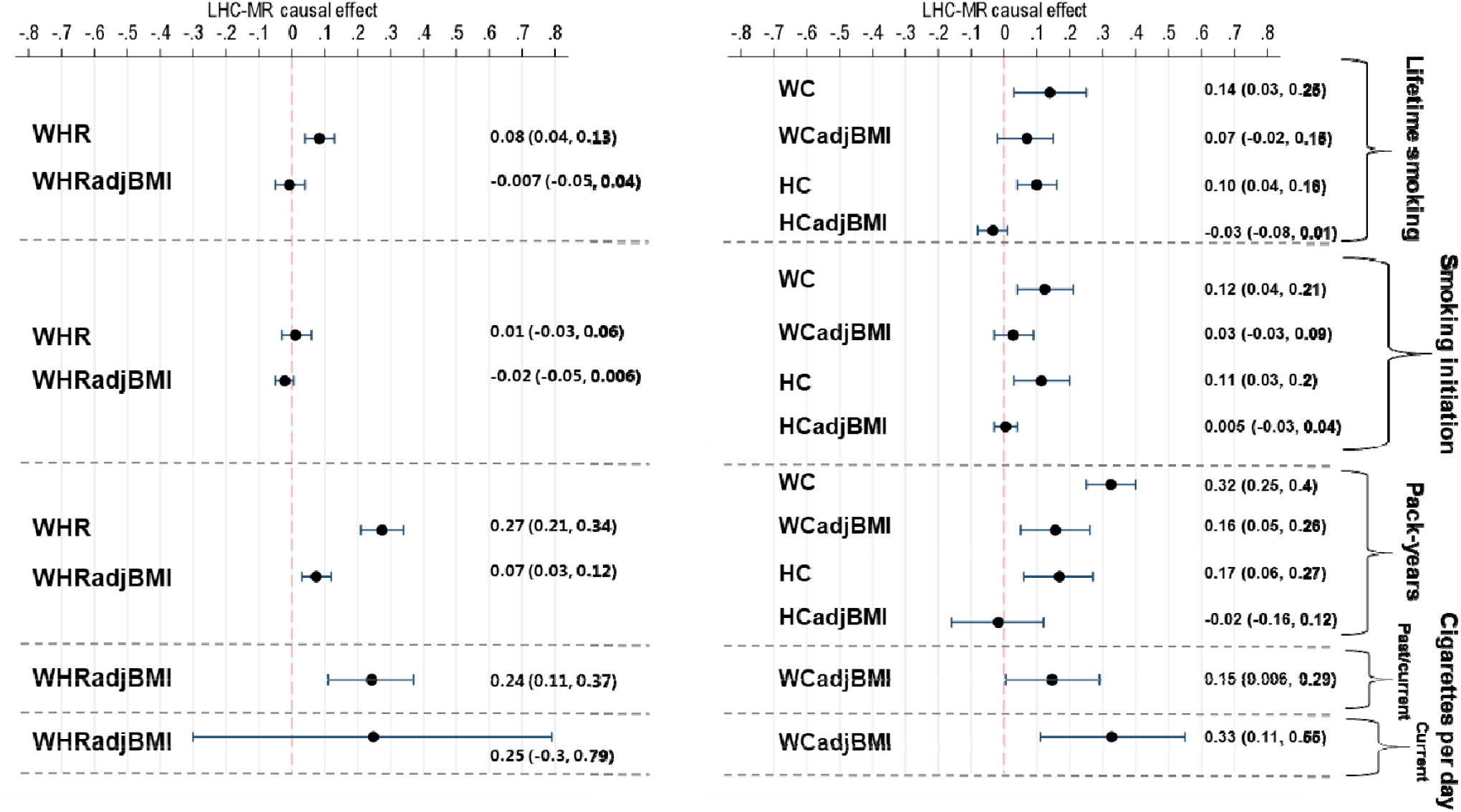
Mendelian randomization results for the causal effect of body fat distribution on smoking traits using the LHC-MR method.

### Association of smoking variants with fat depots and hormonal levels

We investigated the effect of smoking variants on VAT, ASAT, and GSAT tissue volumes by genetic risk score analyses. We generated the scores using the smoking initiation and lifetime smoking variants that showed a causal association with abdominal adiposity in the inverse variance-weighted model. Our results indicated that the genetic risk scores for both smoking initiation and lifetime smoking were associated with a greater increase in VAT compared to ASAT and GSAT. This was demonstrated by the significant association of the genetic scores with VAT/ASAT ratio (0.10 [0.02, 0.17] and 0.25 [0.08, 0.42], respectively) and VAT/GSAT ratio (0.10 [0.03, 0.17] and 0.22 [0.05, 0.39]) (**Figure 5, Supplementary Table 11**). Our findings suggest that the causal effect of smoking initiation and lifetime smoking on WHR is characterized by a disproportionate increase in VAT relative to ASAT and GSAT.

**Figure 5:**
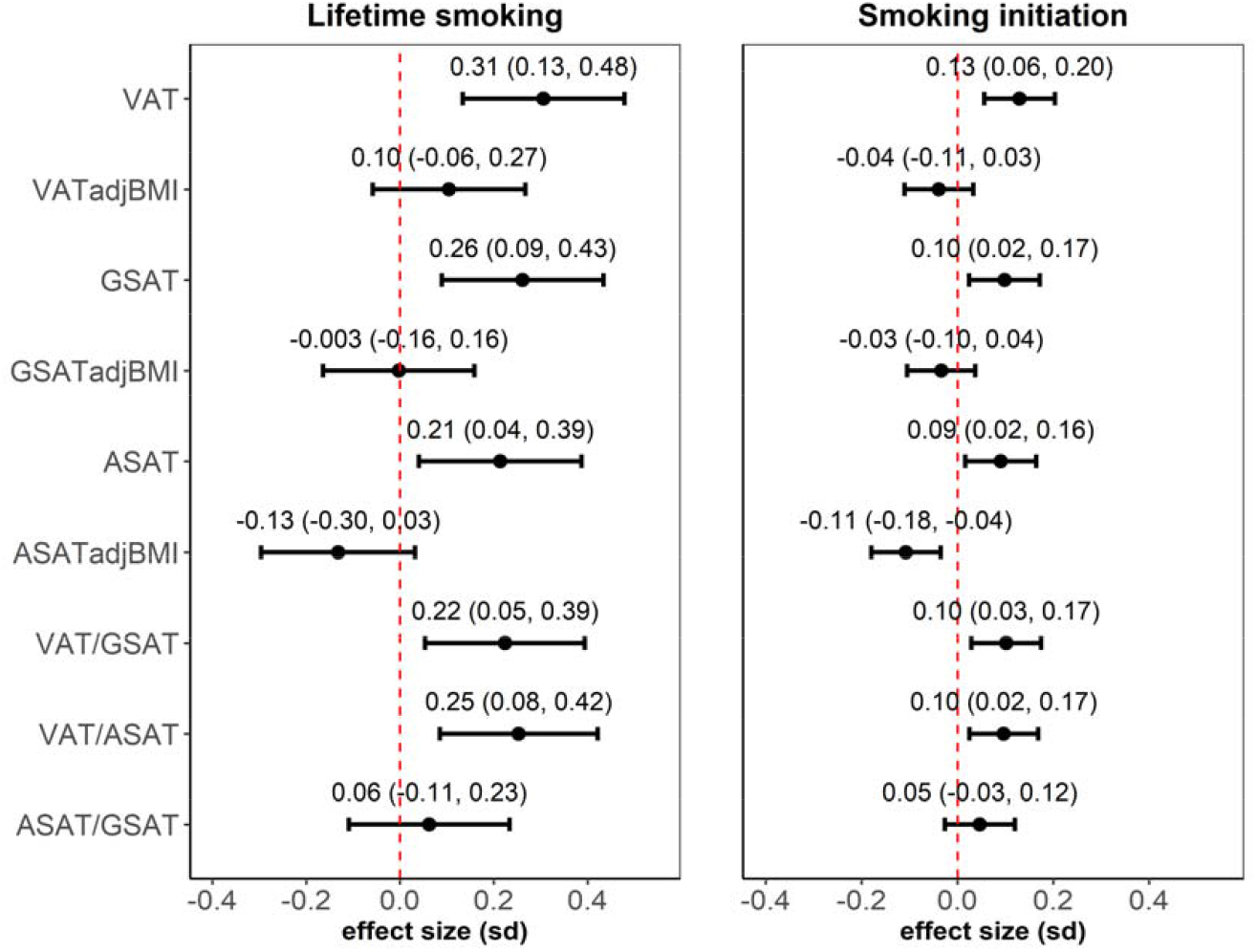
Association of genetic risk scores for lifetime smoking and smoking initiation with visceral adipose tissue (VAT), gluteofemoral fat (GSAT), and abdominal subcutaneous adipose tissue (ASAT) volumes, with and without adjustment for BMI (adjBMI). The associations of the genetic risk scores with VAT/GSAT, VAT/ASAT and ASAT/GSAT ratios are also shown. We constructed genetic risk scores using variants from the inverse variance-weighted model for the causal association between smoking initiation and WHR (121 variants), smoking initiation and WHR_adjBMI_ (130 variants), lifetime smoking and WHR (76 variants), and lifetime smoking and WHR_adjBMI_ (83 variants). The smoking variants causally associated with WHR were used to study associations with BMI-unadjusted fat depots, while the smoking variants causally associated with WHR_adjBMI_ were used to study associations with BMI-adjusted fat depots. We computed weighted genetic scores using the beta of the variants for a smoking trait as weights with gtx.package’s (v0.0.8) grs.summary function. This function approximates the results of a genetic risk score using GWAS summary statistics by calculating the joint effect of genetic variants on an outcome trait.

It has been suggested that smoking may affect abdominal obesity through an impact on hormonal levels, such as cortisol or sex hormones [42, 52-54]. To investigate this, we assessed the effect of the smoking genetic risk scores on the levels of cortisol, oestradiol, testosterone, and sex hormone-binding globulin. The genetic risk scores were not significantly associated with hormonal levels, except for a weak negative association between the score for smoking initiation and sex hormone-binding globulin levels (beta −0.02, 95% CI −0.03 to −0.01) (**Supplementary Figure 5 and Supplementary Table 12**).

## DISCUSSION

Our study showed that smoking initiation and lifetime smoking may causally increase abdominal adipsoity as indicated by higher WHR and WHR_adjBMI_. Analyses for fat depot volumes indicated that visceral fat increases relatively more than abdominal subcutaneous fat. While we found no evidence of an association between smoking heaviness and abdominal fat distribution, our reverse causal analysis indicated that higher abdominal adiposity may causally increase smoking heaviness.

Our findings are in agreement with cross-sectional observational studies that have shown higher abdominal adiposity in smokers than non-smokers [9, 41-44]. However, they do not support a causal relationship between a higher number of cigarettes per day and abdominal adiposity [9, 41, 45]. Previously, three Mendelian randomization studies have used a single variant in the *CHRNA3/5* locus to assess causality between smoking heaviness and abdominal adiposity [14-16]. Two studies found no evidence of causality whereas the third and largest study did [15]. When we examined the Wald ratio estimates for the *CHRNA3/5* locus alone (**Supplementary Table 10**), we found similar estimates supporting a causal relationship between smoking heaviness and abdominal adiposity. However, when we instrumented smoking heaviness using all currently known genetic loci, there was found no evidence of a causal association between smoking heaviness and abdominal adiposity. This suggests that the previous findings based on the *CHRNA3/5* locus alone may have been due to a locus-specific pleiotropic effect on abdominal fat [46-49].

Smoking may increase abdominal fat by increasing either visceral fat or abdominal subcutaneous fat. Our genetic association findings for MRI-based adipose depot volumes suggested that the effect of smoking on abdominal adiposity is mainly driven by an increase in VAT and less the increase in ASAT. These results are in agreement with previous observational studies that have shown a positive association between smoking and higher levels of VAT compared to non-smokers [6, 50]. As the amount of VAT is closely connected to the development of cardiometabolic disease, the observed changes are likely to increase cardiometabolic risk.

A previous Mendelian randomization study reported that BMI may causally increase the likelihood of becoming a smoker and the heaviness of smoking [51]. However, the study did not examine whether there is a causal relationship between abdominal fat distribution and smoking, independent of BMI. While we found no evidence of causality between WHR_adjBMI_ and smoking initiation or lifetime smoking, a higher WHR_adjBMI_ appeared to causally increase the number of cigarettes smoked per day and the total number of pack-years smoked in adulthood. Thus, our results complement previous Mendelian randomization findings by suggesting that body fat distribution may influence smoking heaviness independent of BMI.

It has been suggested that smoking can affect body fat distribution through changes in cortisol and sex hormone levels [42, 52-54]. Smokers have increased levels of cortisol which is linked to insulin resistance and abdominal fat [55]. Smokers have also increased levels oestradiol, testosterone, and sex hormone-binding globulin, which may influence abdominal fat. However, our genetic association results did not support a role of either cortisol or sex hormones in the relationship between smoking and abdominal adiposity.

The strengths of the present study include the use of the largest available GWAS summary-level data on smoking and body fat distribution, and the application of several complementary MR methods and sensitivity analyses to control for pleiotropic effects, sample overlap, latent heritable confounders, reverse causality, and type I error. However, there are also some limitations to our study. Despite performing several sensitivity analyses, we cannot completely rule out the potential influence of residual pleiotropy on our causal estimates. Furthermore, the sample size for body fat distribution in current smokers was relatively small, limiting the statistical power of our smoking heaviness analysis compared to the larger sample sizes used for smoking initiation and lifetime smoking. Finally, our population was restricted to individuals of European genetic ancestry, and thus the findings may not be generalizable to other populations.

In conclusion, smoking initiation and lifetime smoking may causally increase abdominal and particularly visceral fat. Thus, public health efforts to prevent and reduce smoking may also help reduce abdominal fat and the risk of related chronic diseases.

## ACKNOWLEDGEMENTS

The present study was conducted independently without any involvement or influence from any of the funding agencies. Novo Nordisk Foundation Center for Basic Metabolic Research is an independent research center at the University of Copenhagen partially funded by an unrestricted donation from the Novo Nordisk Foundation (NNF18CC0034900). Germán D. Carrasquilla was supported by a grant from the Danish Diabetes Academy that is funded by the Novo Nordisk Foundation (NNF17SA0031406), and from the European Union’s Horizon 2020 research and innovation programme under the Marie Sklodowska-Curie grant agreement No 846502. Tuomas O. Kilpeläinen was supported by the grants NNF17OC0026848, NNF21SA0072102, and NNF22OC0074128 from the Novo Nordisk Foundation.

