## supplementary for "Estimating causality between smoking and abdominal obesity by Mendelian randomization"

**Supplementary note**

**Mendelian randomization using the CAUSE Method**

We implemented the CAUSE Mendelian randomization analysis applying the following analysis steps, previously described in [1-2]:

1. We merged the exposure and outcome summary statistics using the gwas_merge function of the CAUSE R package [1]. We aligned effect sizes of the variants on the exposure and outcome traits to the same allele.
2. We calculated the nuisance parameters to correct for sample overlap between the exposure and outcome GWAS.
3. We clumped the variants using the LDshrink tool with the 1000 Genomes CEU population data as the reference panel.
4. We assigned the priors for the three model parameters – causal effect, shared effect and q. The priors for the causal effect and the shared effect are automatically set to 0 by CAUSE, while for q, the priors are defined by the user. We set the q priors to qalpha = 1 and qbeta = 2.
5. We calculated two models: the sharing model, where the causal effect is set to 0, and the causal model, where posterior probability for the causal effect is calculated.
6. We compared the causal and shared models against the null and against each other, using the expected log pointwise posterior density (ELPD) method to identify which model was most fitting for the data.

**Mendelian randomization using the LHC-MR method**

We implemented the LHC-MR Mendelian randomization method [3] using the following parameters:

1. We filtered the GWAS summary data for exposure and outcome traits by excluding variants with imputation quality score INFO < 0.7 or minor allele frequency (MAF) < 0.01.
2. We excluded variants located in the Major Histocompatibility Complex (MHC), present in chromosome 6 between the base pairs 26M and 34M in human genome build 37.
3. We merged the exposure and outcome GWAS summary statistics with two files containing LD information: 1) LD scores for 4,650,107 common high quality variants with imputation quality score >= 0.99 in the UK Biobank GWAS of Neale Lab and MAF > 0.01 in UK10K and UK Biobank; 2) The local LD distribution of the variants.
4. We assigned the starting points for the parameters that LHC-MR will use to run maximum likelihood estimation (MLE). These are: heritability of trait A, heritability of trait B, confounder effect on trait A, confounder effect on trait B, causal effect of A to B, causal effect of B to A, and the intercept of the genetic correlation between A and B.
5. We performed the maximum likelihood estimation.

**Mendelian randomization using the inverse variance-weighted (IVW), MR-Egger, weighted median, and weighted mode methods**

For the inverse variance-weighted, MR-Egger, weighted median, and weighted mode Mendelian randomization methods, we applied a range of sensitivity tests (**Supplementary Table 6**) and plots (**Supplementary Figure 3**) to assess pleiotropy and ensure that the reported causal effects were unbiased. We considered the results unbiased if they met the following criteria:

1. The causal estimates were considered heterogeneous if the p-value for Cochran’s Q test was <0.05 and I^2^ was >0.25.
2. Egger’s intercept with a p-value <0.05 was considered as evidence that the results are likely to be affected by horizontal pleiotropy. We interpreted the results of Egger’s intercept test together with Rucker test results. Rucker test compares Cochran’s Q calculated with IVW and MR-Egger causal effects and reports which method is a better fit for the data: If the Cochran’s Q is significantly different between IVW and MR-Egger (Rucker test p-value < 0.05), then MR-Egger, which controls for horizontal pleiotropy, is considered to be a better fit for the data.
3. We visualized the effects of heterogeneity and horizontal pleiotropy using leave-one-out forest plots and funnel plots (**Supplementary Figure 3**). If these sensitivity plots showed evidence of outlier variants, we used PhenoScanner [4-5] to examine whether the outlier variants could be possible confounders and were strongly associated with other traits than the smoking-abdominal adiposity relationship.

If the sensitivity tests and plots and PhenoScanner results suggested the presence of horizontal pleiotropy, we implemented the following additional steps:

1. We detected pleiotropic variants using RadialMR’s iterative Cochran’s Q method (P threshold <0.05). The iterative Cochran’s Q, either IVW’s Q or Egger’s Q’, was chosen depending on the result from the Rucker framework test.
2. We excluded any pleiotropic variants and re-implemented the Mendelian Randomization analyses without including these variants. We then reapplied the sensitivity tests and plots to assess whether we had successfully removed the variants introducing horizontal pleiotropy.

**Interpretation of the results indicative of causality for inversed variant weighted (IVW), MR-Egger, weighted median, and weighted mode methods:**

In the following, we describe the results of the sensitivity analyses and the interpretation of the results of the Mendelian randomization analyses using the IVW, MR-Egger, weighted median, and weighted mode methods.

*Lifetime smoking → Abdominal adiposity*

The IVW and the weighted median Mendelian randomization methods showed a positive causal association of lifetime smoking with WHR and WC_adjBMI_. The IVW method also showed a positive causal association of lifetime smoking with WHR_adjBMI_. (**Figure 1-2 and** **Supplementary Table 5**). The Cochran’s Q test (P > 0.05) and I^2^ value (<25%) showed no evidence of heterogeneity between the causal estimates of the instrumental variables. Furthermore, Egger’s intercept was not significant and Rucker’s test suggested that IVW is a better fit for the data than MR-Egger (**Supplementary Table 5**). However, RadialMR and the sensitivity plots identified five potentially pleiotropic variants for lifetime smoking**→**WHR_adjBMI_ (rs2867112, rs4543592, rs6778080, rs8078228 and rs889398) and three for lifetime smoking**→**WC_adjBMI_ (rs1050847, rs17309874 and rs6778080). When checking these variants in PhenoScanner, we did not detect any clear important confounder. Yet, the variant rs6778080 was associated with age at menarche and blood count essay parameters, and rs8078228 with asthma and white cell count.Therefore, for lifetime smoking**→**WHR_adjBMI_ we report the Mendelian randomization results after removing the five pleiotropic variants. After the extraction of the outliers, the sensitivity tests and sensitivity plots showed no evidence of pleiotropy (**Supplementary Table 6, Supplementary Figure 2 and 3**). Overall, our findings supported a positive causal association between lifetime smoking and WHR, WHR_adjBMI_ and WC_adjBMI._

*Smoking initiation → Abdominal adiposity*

The IVW and weighted median Mendelian randomization methods showed a positive causal association of smoking initiation with WHR and WC, and the weighted mode method also showed a positive causal association of smoking initiation with WHR (**Figure 1-2 and** **Supplementary Table 5**). The Cochran’s Q (P > 0.05) and the I^2^ value (<25%) did not suggest heterogeneity between the causal estimates of the instrumental variables. Furthermore, Egger’s intercept was not significant and Rucker test indicated that IVW was a better fit with the data than MR-Egger (**Supplementary Table 5**). The sensitivity plots identified the variant rs1349979 as a potential pleiotropic variant (**Supplementary Figure 2 and 3)** and PhenoScanner did not detect any potential confounder. Overall, our findings support a positive causal association between smoking initiation and WHR and WC.

*Pack-years → Abdominal adiposity*

The weighted median and weighted mode Mendelian randomization methods showed a negative causal association of pack-years of smoking with HC. The IVW method showed a positive causal association of pack-years of smoking with WHR_adjBMI_ and a negative causal association with WC. (**Figure 1-2 and** **Supplementary Table 5**). The Cochran’s Q test was not significant (P >= 0.05), whereas the I^2^ value showed evidence of heterogeneity (>25%) for pack-years **→** WC and pack-years **→** HC. The Egger’s intercept was significant and tge Rucker test indicated that the MR-Egger method has a bitter fit with the data than IVW for pack-years **→** WC and pack-years **→** HC (**Supplementary Table 5**). The sensitivity plots and RadialMR identified one potentially pleiotropic variant for pack-years **→** WHR_adjBMI_ (rs10519203) and two variants for pack-years **→** WC and pack-years **→** HC (rs136403 and rs150353) (**Supplementary Figure 2 and 3)**. None of the aforementioned pleiotropic variants showed strong associations with potential confounders in PhenoScanner. We reported the Mendelian results for pack-years **→** WC and pack-years **→** HC after removing each of the potential pleiotropic variants. After the outlier extraction, the sensitivity tests and sensitivity plots suggested remaining influence of pleiotropy on the results for pack-years **→** HC (**Supplementary Table 6, Supplementary Figure 2 and 3**). Thus, the Mendelian randomization results do not provide robust support for a causal effect of pack-years of smoking on abdominal adiposity.

*Cigarettes per day (past/current) → Abdominal adiposity*

None of the Mendelian randomization methods showed a significant causal association of cigarettes per day (past/current) with WHR_adjBMI_ or WC_adjBMI_, even though each of the methods suggested a positive direction of the causal estimate (**Figure 1-2 and** **Supplementary Table 5).** The Cochran’s Q test (P > 0.05) and I^2^ value (<25%) indicated an absence of strong heterogeneity between the causal estimates of the instrumental variables. Furthermore, Egger’s intercept was non-significant, and Rucker test indicated that IVW shows a better fit with the data than MR-Egger (**Supplementary Table 5**). The sensitivity plots were consistent with the sensitivity tests (**Supplementary Figure 2 and 3)**. Thus, our findings suggest no causal association between cigarettes per day (past/current) and abdominal adiposity.

*Cigarettes per day (current) → Abdominal adiposity*

The IVW and weighted median Mendelian randomization methods showed a positive causal association of cigarettes per day (current) with WHR_adjBMI_. The Cochran’s Q test (P > 0.05) and I^2^ value (<25%) showed no evidence of heterogeneity between the causal estimates of the instrumental variables. Furthermore, Egger’s intercept was non-significant, and Rucker test indicated that IVW is a better with the data than MR-Egger (**Supplementary Table 5**). However, the funnel plot suggested that when removing the rs1317286 variant, the causal effect in the IVW model was no longer present (**Supplementary Figure 2 and 3)**. To note, rs1317286 represents the strongest known smoking heaviness-locus in CHRNA3/5 and PhenoScanner showed strong associated with smoking heaviness traits. Thus, we further test the locus-specific pleiotropic effects on discrete traits for the variant rs1317286 (Supplementary Figure 4). Thus, our findings suggest that the positive causal association between cigarettes per day (current) and WHR_adjBMI_ may be exclusively driven by the single variant rs1317286.

**Multivariable Mendelian randomization using the MVMR package**

We used the MVMR package, following the steps previously described in [6], with the exception that we used the function qhet_mvmr that takes pleiotropy into account when estimating the causal effects. Since the ghet_mvmr function is still under development, we also assessed whether pleiotropy may affect the results by manually removing the outliers (step 8 below).

1. We identified independent variants by obtaining genome-wide significant variants for each risk factor, and clumping them using the ld_clump_local function in the package ieugwasr, using the updated European 1000 Genomes reference panel with the following settings: clump_kb = 10000kb and clump_r^2^ < 0.001.
2. We replaced the independent lead variants not present in the outcome trait GWAS summary statistics by proxy variants (r^2^ > 0.8 in European 1000 Genomes reference panel). To identify the most accurate proxy, we clumped (clump_kb = 10000kb and clump_r^2^ < 0.001) all proxies found in the locus of the missing lead variant, using the p-value for the exposure trait for which the missing lead variant was genome-wide significant. If the missing lead variant was found to be genome-wide significant for more than one trait, the p-value of the exposure trait for which the missing variant presented the lowest p-value was used.
3. We harmonized the risk factor and outcome data, so that for each variant, all the genetic effect sizes were aligned to the same allele, while removing palindromic variants.
4. For each SNP, we calculated a covariance matrix using the function phenocov_mvmr(). This function calculates the covariance between the risk factors for each variant using the standard error for each risk factor and the phenotypic correlation between the risk factors in the form of a phenotypic correlation matrix. In this case, the phenotypic correlation between lifetime smoking and education was set to -0.15. The reason for using this correlation cut-off was that the Spearman correlation coefficient in the UK Biobank for “lifetime smoking”, smoking initiation, and packs per year vs. education was -0.1392, -0.0904, and -0.1837, respectively (all with p < 0.00001).
5. We calculated the conditional F-statistic of the set of independent variants with each risk factor. The conditional F-statistic was calculated using the function strength_mvmr that calculates the analog of the F-statistic in a multivariable setting. If the conditional F-statistic for lifetime smoking or for education was < 10, we considered that the set of independent variants was weakly associated with the risk factors.
6. If the conditional F-statistic for all risk factors was > 10, we calculated conditional Cochran’s Q to test whether the causal effect estimated with IVW may be biased due to pleiotropy. We considered a p-value < 0.05 in the conditional Cochran’s Q test as evidence of pleiotropy.
7. We calculated IVW causal effects using the ivw_mvmr() function.
8. To test whether the IVW causal effect may be biased by pleiotropy, we re-implemented steps 4-7 after removing the outlier variants that could introduce pleiotropy. We calculated the conditional Cochran’s Q for each variant and removed variants with Cochran’s Q > 3.84. While this threshold is stringent and may have removed variants that were not true outliers, we used this to ensure that our results hold without outlier variants.

**Supplementary Note References:**

1. Jean Morrison (2020). CAUSE software tutorial. Retrieved from <https://jean997.github.io/cause/ldl_cad.html>
2. Morrison J, Knoblauch N, Marcus JH, Stephens M, He X. Mendelian randomization accounting for correlated and uncorrelated pleiotropic effects using genome-wide summary statistics. Nat Genet. 2020;52(7):740-7. Epub 2020/05/27. doi: 10.1038/s41588-020-0631-4. PubMed PMID: 32451458; PubMed Central PMCID: PMCPMC7343608.
3. Darrous L, Ninon M, and Zoltán K. Simultaneous estimation of bi-directional causal effects and heritable confounding from GWAS summary statistics. Nature communications 12.1 (2021): 1-15.
4. James R Staley, et al. PhenoScanner: a database of human genotype-phenotype associations. Bioinformatics 2016; 32(20):3207-3209
5. Mihir A Kamat, et al. PhenoScanner V2: an expanded tool for searching human genotype-phenotype associations. Bioinformatics 2019; 35(22):4851-4853
6. Sanderson E, Davey Smith G, Windmeijer F, Bowden J. An examination of multivariable Mendelian randomization in the single-sample and two-sample summary data settings. Int J Epidemiol. 2019;48(3):713-27. Epub 2018/12/12. doi: 10.1093/ije/dyy262. PubMed PMID: 30535378; PubMed Central PMCID: PMCPMC6734942.
7. Sanderson E, Spiller W, Bowden J. Testing and correcting for weak and pleiotropic instruments in two-sample multivariable mendelian randomisation. Biorxiv. 2020

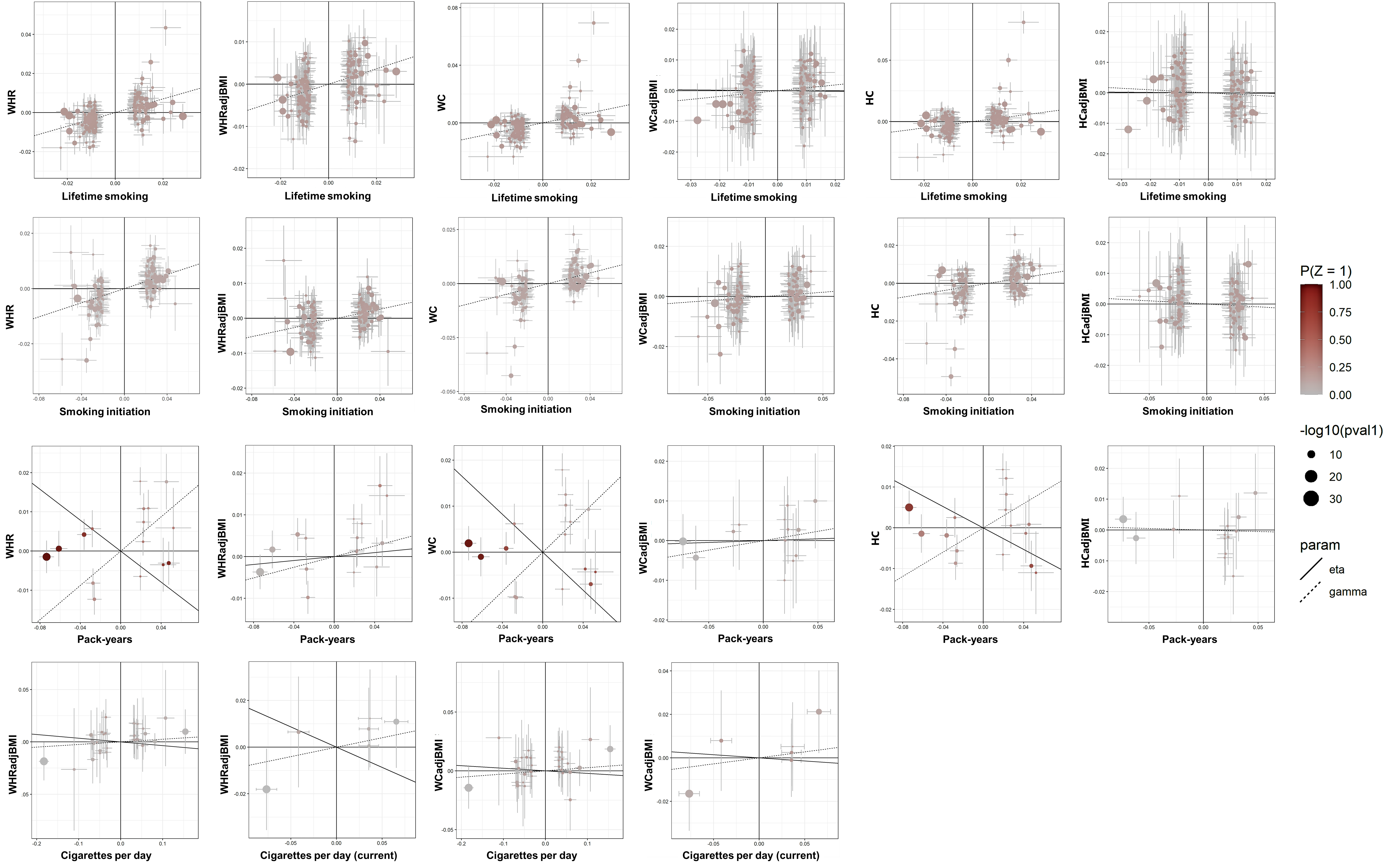

**Supplementary Figure 1.** Scatter plots showing the effect estimates for smoking traits (x-axis) plotted against the estimates for body fat distribution traits (y-axis) for the causal model using the CAUSE method. The size of the circles indicates the P value of the association of the variant with the smoking trait and the color indicates the P value for the association with the body fat distribution trait. Only variants with P<5x10^−8^ for association with the smoking trait are included in the plot.

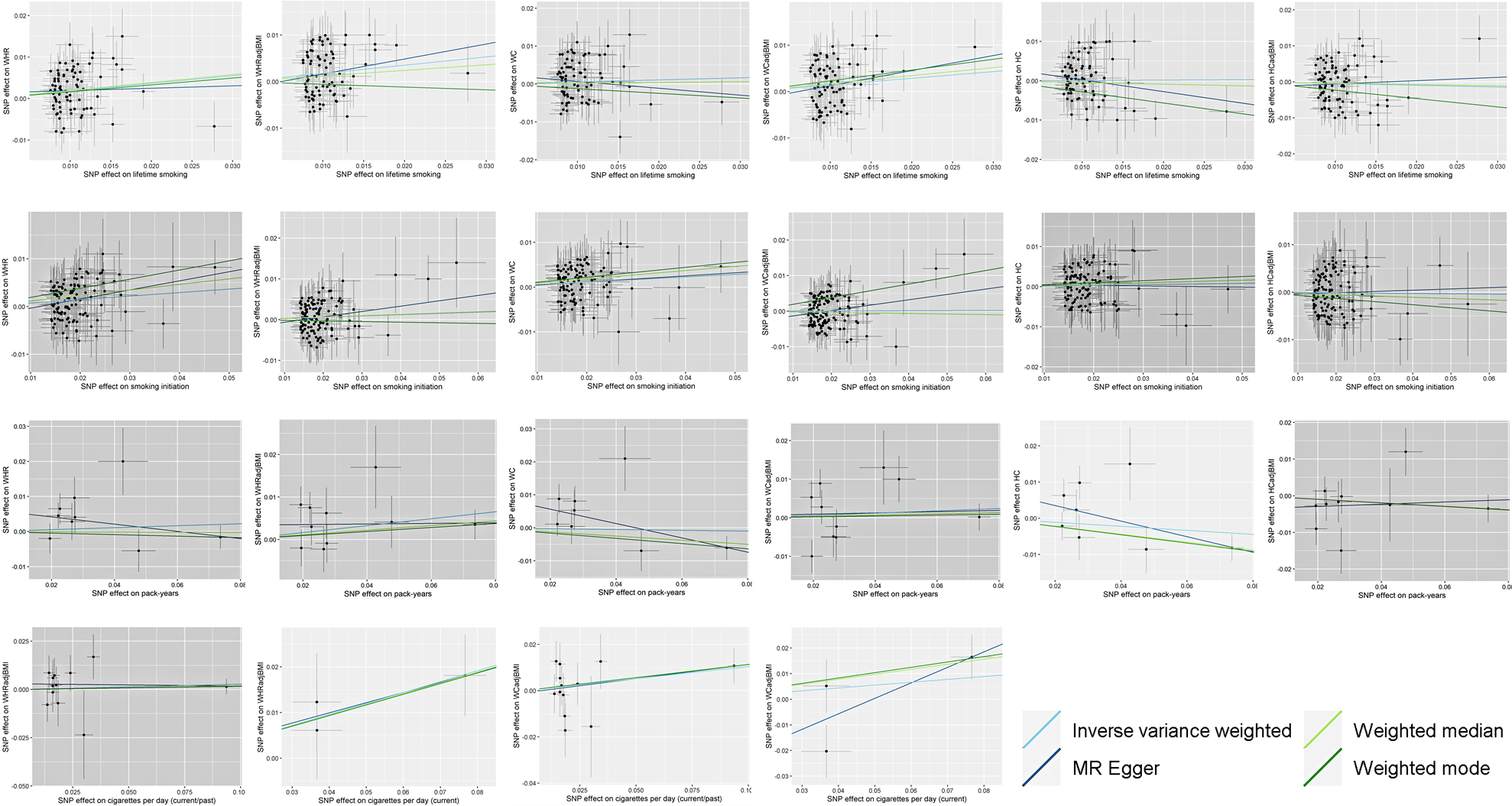

**Supplementary Figure 2.** Scatter plots showing the effect estimates for smoking traits (x-axis) plotted against the estimates for body fat distribution traits (y-axis) in the IVW, MR-Egger, weighted median and weighted mode methods after outlier removal and accounting for pleiotropy.

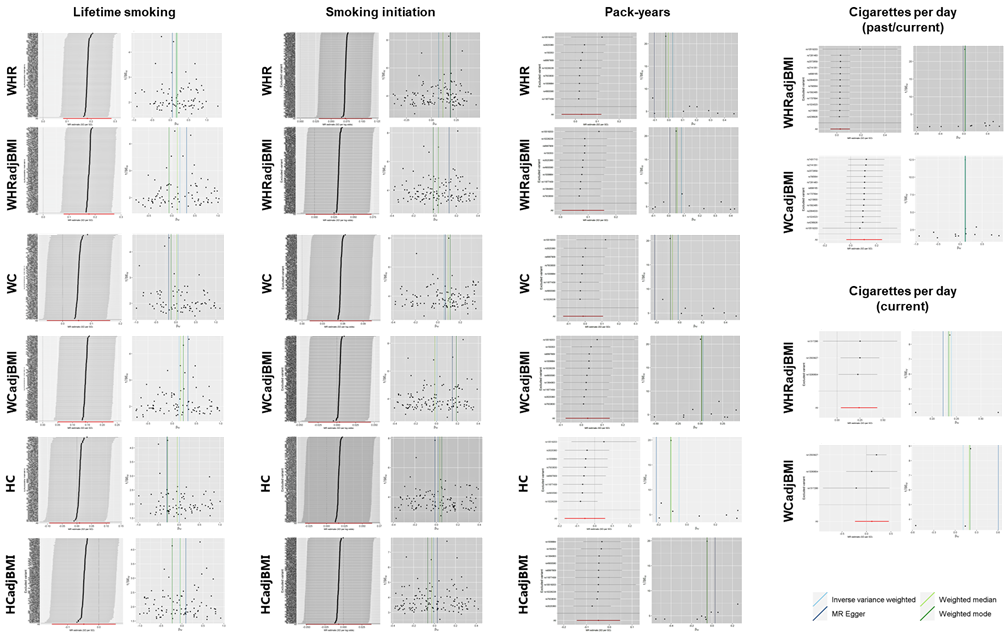

**Supplementary Figure 3.** Leave-one-out and funnel sensitivity plots for the inverse variance-weighted, MR-Egger, weighted median and weighted mode methods.

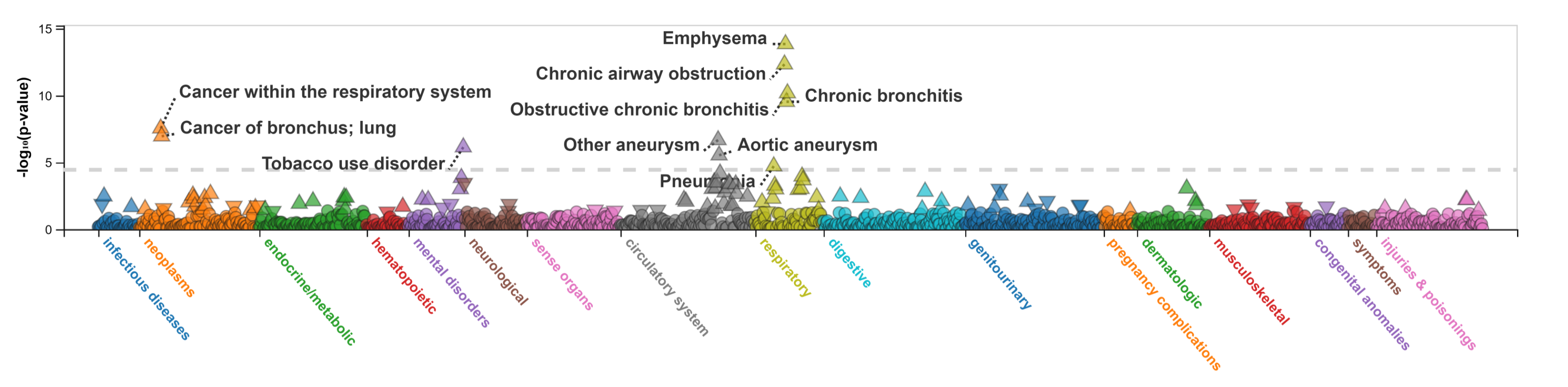

**Supplementary Figure 4.** Phenome-wide association analysis for the rs1317286 (*CHRNA3/5*) variant in the UK Biobank including over 400,000 European-ancestry individuals. Source: <http://pheweb.sph.umich.edu/>

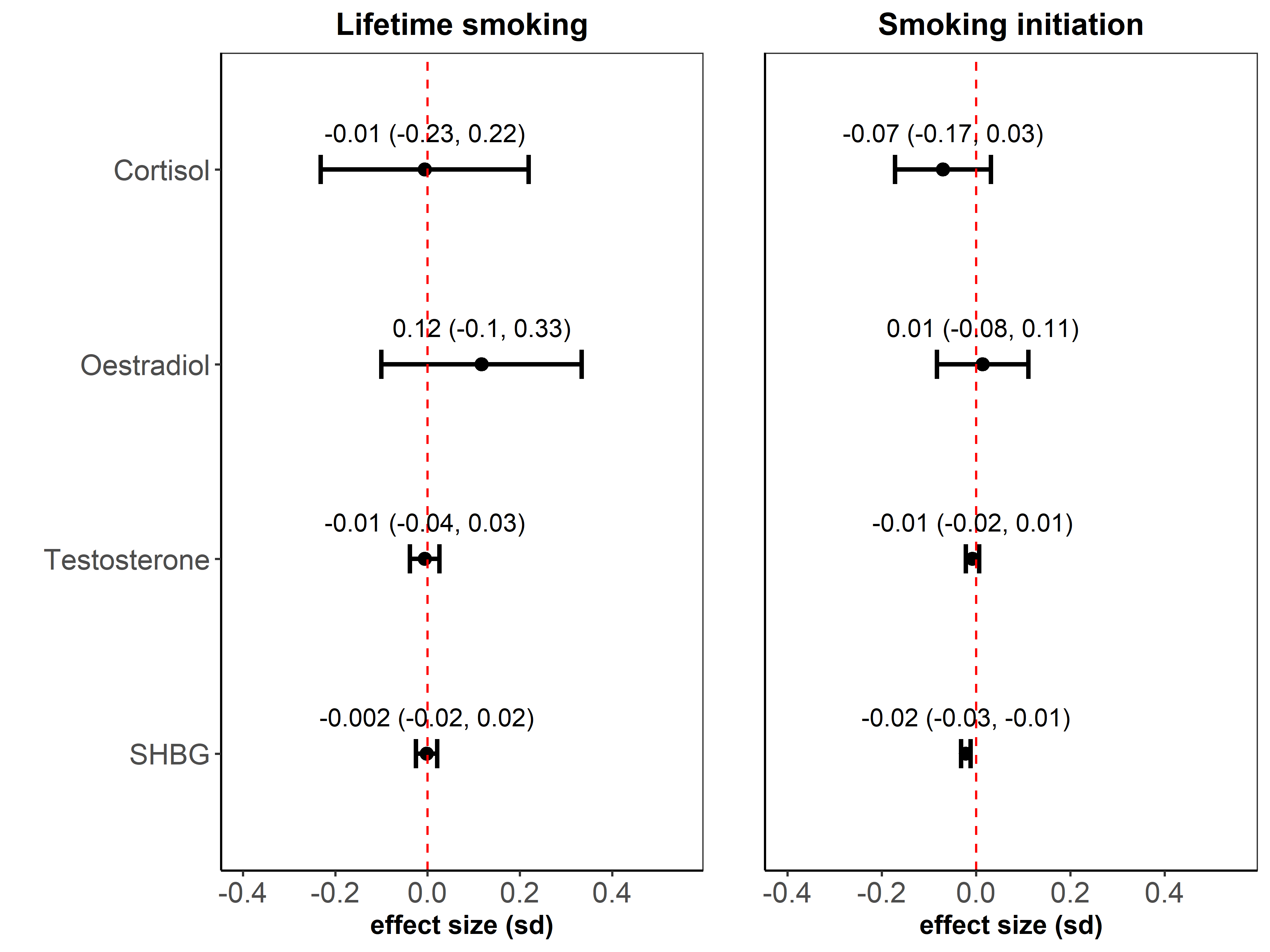

**Supplementary Figure 5.** Association of genetic risk scores for lifetime smoking and smoking initiation with hormonal levels, including cortisol, testosterone, oestradiol and sex hormone-binding globulin (SHBG). We constructed genetic risk scores using variants from the inverse variance-weighted model for the causal association between smoking initiation and WHR_adjBMI_ (130 variants) and lifetime smoking and WHR_adjBMI_ (83 variants). We computed weighted genetic scores using the beta of the variants for the smoking trait as weights with gtx.package’s (v0.0.8) grs.summary function. This function approximates the results of a genetic risk score using GWAS summary statistics by calculating the joint effect of genetic variants on an outcome trait.

| **Supplementary Table 1: Datasets for smoking and body fat distribution traits used in the present study** | | | |
| --- | --- | --- | --- |
| **Trait and MR Method** | **Author** | **Sample size** | **Population/note** |
| **Lifetime smoking** |  |  |  |
| CAUSE, LHC-MR, 2SMR and MVMR | Wootton et al 2020 | 462,690 | European individuals from UK Biobank |
| **Smoking Initiation** |  |  |  |
| CAUSE, LHC-MR, and MVMR | Liu et al 2019 | 632,802 | European individuals from GSGC-UKBB meta-analysis without 23andMe. |
| 2SMR | Liu et al 2019 | 1,232,091 | European individuals from GSGC-UKBB meta-analysis with 23andMe. |
| **Pack-years** |  |  |  |
| CAUSE and LHC-MR | Elseworth et al 2018 | 142,387 | European individuals from UK Biobank |
| 2SMR | Elseworth et al 2019 | 142,387 | European individuals from UK Biobank |
| **Cigarettes per day (past and current)** |  |  |  |
| CAUSE and LHC-MR | Liu et al 2019 | 263,954 | European individuals from GSGC-UKBB meta-analysis without 23andMe. |
| 2SMR | Liu et al 2019 | 337,334 | European individuals from GSGC-UKBB meta-analysis with 23andMe. |
| **Cigarettes per day (current)** |  |  |  |
| CAUSE and LHC-MR | Elseworth et al 2018 | 33,229 | European individuals from UK Biobank |
| 2SMR | Elseworth et al 2018 | 33,229 | European individuals from UK Biobank |
| **WHR** |  |  |  |
| CAUSE and LHC-MR | Pulit et al 2019 | 697,734 | European individuals from GIANT-UK Biobank meta-analysis |
| 2SMR and MVMR | Shungin et al 2015 | 212,248 | European individuals from GIANT |
| **WHRadjBMI** |  |  |  |
| CAUSE and LHC-MR | Pulit et al 2019 | 694,649 | European individuals from GIANT-UK Biobank meta-analysis |
| 2SMR and MVMR | Shungin et al 2015 | 210,086 | European individuals from GIANT. Used for lifetime smoking and smoking initiation. |
| **WC** |  |  |  |
| CAUSE and LHC-MR | Elseworth et al 2018 | 462,166 | European individuals from UKBB |
| 2SMR and MVMR | Shungin et al 2015 | 232,101 | European individuals from GIANT |
| **WCadjBMI** |  |  |  |
| CAUSE, LHC-MR, 2SMR and MVMR | Shungin et al 2015 | 231,355 | European individuals from UKBB |
| **WHRadjBMI stratified for smokers** |  |  |  |
| CAUSE and LHC-MR | Justice et al 2017 | 40,543 | European individuals. Used for cigarettes per day analysis |
| 2SMR | Justice et al 2017 | 40,543 | European individuals. Used for cigarettes per day analysis |
| **WCadjBMI stratified for smokers** |  |  |  |
| CAUSE and LHC-MR | Justice et al 2017 | 43,226 | European individuals. Used for cigarettes per day analysis |
| 2SMR | Justice et al 2017 | 43,226 | European individuals. Used for cigarettes per day analysis |
| **HC** |  |  |  |
| CAUSE and LHC-MR | Elseworth et al 2018 | 462,117 | European individuals from UKBB |
| 2SMR and MVMR | Shungin et al 2015 | 213,038 | European individuals from GIANT |
| **HCadjBMI** |  |  |  |
| CAUSE and LHC-MR | Shungin et al 2015 | 211,117 | European individuals from GIANT |
| 2SMR and MVMR | Shungin et al 2015 | 211,117 | European individuals from GIANT. Used for lifetime smoking and smoking initiation. |
| CAUSE, Causal Analysis Using Summary Effect Estimates; 2SMR, two-sample Mendelian randomization; MVMR, multivariable Mendelian randomization; LHC-MR, Latent Heritable Confounder Mendelian randomization; WHR, waist-hip ratio; WHRAdjBMI, waist-hip ratio adjusted for body mass index; WC, waist circumference; WCadjBMI, waist circumference adjusted for body mass index; HC, hip circumference; HCadjBMI, hip circumference adjusted for body mass index. | | | |

| **Supplementary Table 2. Results for Mendelian randomization analyses using the CAUSE method** | | | | | | | | | |
| --- | --- | --- | --- | --- | --- | --- | --- | --- | --- |
|  | **Lifetime smoking** | **Smoking initiation** | **Pack-years** |  | **Lifetime smoking** | **Smoking initiation** | **Pack-years** | **Cigarettes per day (past/current)** | **Cigarettes per day (current)** |
|  | **WHR** | | |  | **WHRadjBMI** | | | | |
| Median causal effect (95% CI) | 0.35 (0.29, 0.41) | 0.13 (0.1, 0.16) | 0.22 (-0.01, 0.3) |  | 0.18 (0.13, 0.24) | 0.07 (0.03, 0.10) | 0.06 (0.01, 0.15) | 0.02 (-0.1, 0.1) | 0.08 (-0.47, 0.27) |
| Median shared effect (95% CI) | 0 (-0.29, 0.27) | 0 (-0.15, 0.16) | -0.2 (-0.28, 0.27) |  | 0 (-0.27, 0.24) | 0 (-0.16, 0.16) | 0.02 (-0.2, 0.31) | -0.04 (-0.7, 0.36) | -0.17 (-1.56, 0.91) |
| Median q (95% CI) | 0.18 (0, 0.86) | 0.18 (0, 0.86) | 0.3 (0.09, 0.83) |  | 0.18 (0, 0.86) | 0.18 (0, 0.86) | 0.19 (0, 0.85) | 0.14 (0, 0.86) | 0.19 (0.01, 0.85) |
| Sharing vs causal *P* | 1.30E-06 | 2.10E-15 | 0.85 |  | 2.20E-09 | 5.20E-03 | 0.13 | 0.68 | 0.58 |
|  | **WC** | | |  | **WCadjBMI** | | | | |
| Median causal effect (95% CI) | 0.35 (0.29, 0.41) | 0.12 (0.09, 0.16) | 0.21 (-0.02, 0.28) |  | 0.09 (-0.05, 0.23) | 0.03 (-0.04, 0.11) | 0.05 (-0.03, 0.15) | 0.03 (-0.11, 0.11) | 0.06 (-0.27, 0.29) |
| Median shared effect (95% CI) | 0 (-0.29, 0.28) | 0 (-0.16, 0.17) | -0.2 (-0.29, 0.27) |  | -0.01 (-0.63, 0.5) | 0 (-0.31, 0.34) | 0.01 (-0.43, 0.46) | -0.02 (-0.48, 0.4) | -0.03 (-0.89, 0.74) |
| Median q (95% CI) | 0.18 (0, 0.86) | 0.18 (0, 0.86) | 0.23 (0.06, 0.86) |  | 0.19 (0, 0.86) | 0.19 (0, 0.84) | 0.17 (0, 0.86) | 0.19 (0, 0.86) | 0.22 (0.01, 0.85) |
| Sharing vs causal *P* | 7.70E-12 | 7.30E-08 | 0.86 |  | 0.15 | 0.26 | 0.25 | 0.71 | 0.91 |
|  | **HC** | | |  | **HCadjBMI** | | | | |
| Median causal effect (95% CI) | 0.26 (0.18, 0.33) | 0.09 (0.05, 0.13) | 0.15 (-0.03, 0.25) |  | -0.05 (-0.23, 0.13) | -0.02 (-0.1, 0.05) | -0.01 (-0.13, 0.11) | NA | NA |
| Median shared effect (95% CI) | 0 (-0.31, 0.31) | 0 (-0.18, 0.19) | -0.13 (-0.29, 0.26) |  | -0.01 (-0.72, 0.63) | 0 (-0.39, 0.36) | 0 (-0.57, 0.6) | NA | NA |
| Median q (95% CI) | 0.19 (0, 0.86) | 0.18 (0, 0.86) | 0.25 (0.02, 0.86) |  | 0.2 (0.01, 0.86) | 0.18 (0, 0.84) | 0.17 (0, 0.87) | NA | NA |
| Sharing vs causal *P* | 5.40E-09 | 1.20E-03 | 0.79 |  | 0.50 | 0.61 | 0.99 | NA | NA |
| The results display the results according to the goodness-of-fit for the causal or the sharing model. The median q value indicates the proportion of variants that showed evidence of correlated pleiotropy. WHR, waist-to-hip ratio; WHRAdjBMI, waist-hip ratio adjusted for body mass index; WC, waist circumference; WCadjBMI, waist circumference adjusted for body mass index; HC, hip circumference; HCadjBMI, hip circumference adjusted for body mass index; CI, confidence interval; NA, not available. | | | | | | | | | |

| **Supplementary Table 3: CAUSE expected log pointwise posterior density (ELPD) results** | | | | | | | | | | | | | | | | | | | | |
| --- | --- | --- | --- | --- | --- | --- | --- | --- | --- | --- | --- | --- | --- | --- | --- | --- | --- | --- | --- | --- |
|  |  |  | **WHR** | | | **WHRadjBMI** | | | **WC** | | | **WCadjBMI** | | | **HC** | | | **HCadjBMI** | | |
| **Smoking trait** | **Model 1** | **Model 2** | **Delta_ELPD** | **SE_delta_ELPD** | **z** | **Delta_ELPD** | **SE_delta_ELPD** | **z** | **Delta_ELPD** | **SE_delta_ELPD** | **z** | **Delta_ELPD** | **SE_delta_ELPD** | **z** | **Delta_ELPD** | **SE_delta_ELPD** | **z** | **Delta_ELPD** | **SE_delta_ELPD** | **z** |
| Lifetime smoking | Null | Sharing | -194.83 | 15.60 | -12.49 | -85.62 | 11.07 | -7.73 | -184.21 | 15.22 | -12.10 | -6.49 | 3.40 | -1.91 | -78.45 | 10.01 | -7.84 | -0.71 | 1.53 | -0.47 |
|  | Null | Causal | -196.28 | 15.80 | -12.42 | -86.88 | 11.27 | -7.71 | -185.65 | 15.42 | -12.04 | -7.08 | 3.94 | -1.80 | -79.78 | 10.23 | -7.80 | -0.71 | 2.06 | -0.35 |
|  | Sharing | Causal | -1.46 | 0.31 | -4.71 | -1.26 | 0.21 | -5.87 | -1.44 | 0.21 | -6.74 | -0.59 | 0.56 | -1.05 | -1.34 | 0.23 | -5.72 | 5.06E-04 | 0.54 | 9.32E-04 |
| Smoking initiation | Null | Sharing | -111.83 | 11.91 | -9.39 | -53.84 | 9.58 | -5.62 | -91.71 | 10.96 | -8.37 | -3.52 | 2.71 | -1.30 | -39.33 | 7.19 | -5.47 | -0.34 | 1.25 | -0.28 |
|  | Null | Causal | -113.20 | 12.07 | -9.38 | -54.83 | 9.87 | -5.55 | -92.94 | 11.18 | -8.31 | -3.86 | 3.20 | -1.21 | -40.68 | 7.63 | -5.33 | -0.20 | 1.77 | -0.11 |
|  | Sharing | Causal | -1.37 | 0.17 | -7.85 | -0.98 | 0.38 | -2.56 | -1.23 | 0.23 | -5.26 | -0.35 | 0.53 | -0.65 | -1.35 | 0.44 | -3.03 | 0.15 | 0.53 | 0.28 |
| Pack-years | Null | Sharing | -35.29 | 8.11 | -4.35 | -8.81 | 3.76 | -2.34 | -44.65 | 8.42 | -5.30 | -0.76 | 1.25 | -0.60 | -20.41 | 5.63 | -3.62 | 0.67 | 0.22 | 3.10 |
|  | Null | Causal | -34.47 | 8.43 | -4.09 | -9.90 | 4.44 | -2.23 | -42.74 | 9.25 | -4.62 | -1.36 | 2.10 | -0.65 | -18.95 | 6.57 | -2.88 | 1.53 | 0.54 | 2.81 |
|  | Sharing | Causal | 0.82 | 0.80 | 1.03 | -1.09 | 0.98 | -1.11 | 1.91 | 1.61 | 1.18 | -0.60 | 0.88 | -0.69 | 1.46 | 1.81 | 0.81 | 0.86 | 0.35 | 2.45 |
| Cigarettes per day (past/current) | Null | Sharing | NA | NA | NA | 0.47 | 0.41 | 1.14 | NA | NA | NA | 0.42 | 0.33 | 1.28 | NA | NA | NA | NA | NA | NA |
|  | Null | Causal | NA | NA | NA | 0.79 | 1.09 | 0.73 | NA | NA | NA | 0.76 | 0.94 | 0.81 | NA | NA | NA | NA | NA | NA |
|  | Sharing | Causal | NA | NA | NA | 0.32 | 0.69 | 0.47 | NA | NA | NA | 0.34 | 0.61 | 0.55 | NA | NA | NA | NA | NA | NA |
| Cigarettes per day (current) | Null | Sharing | NA | NA | NA | 0.50 | 0.38 | 1.34 | NA | NA | NA | 0.44 | 0.12 | 3.74 | NA | NA | NA | NA | NA | NA |
|  | Null | Causal | NA | NA | NA | 0.70 | 1.28 | 0.55 | NA | NA | NA | 1.01 | 0.54 | 1.87 | NA | NA | NA | NA | NA | NA |
|  | Sharing | Causal | NA | NA | NA | 0.19 | 0.92 | 0.21 | NA | NA | NA | 0.57 | 0.43 | 1.32 | NA | NA | NA | NA | NA | NA |
| WHR, waist-hip ratio; WHRAdjBMI, waist-hip ratio adjusted for body mass index; WC, waist circumference; WCadjBMI, waist circumference adjusted for body mass index; HC, hip circumference; HCadjBMI, hip circumference adjusted for body mass index; SE, standard error; NA, not available. | | | | | | | | | | | | | | | | | | | | |

| **Supplementary Table 4: LHC-MR causal effects of heritable shared confounders on smoking and body composition traits** | | | | | | |
| --- | --- | --- | --- | --- | --- | --- |
|  | **Confounder term for smoking** | | | **Confounder term for adiposity** | | |
| **Exposure-outcome pairs** | **Coefficient** | **SE** | **P-value** | **Coefficient** | **SE** | **P-value** |
| Lifetime smoking-WHR | 0,09 | 0,01 | 2,91E-13 | -0,08 | 0,02 | 2,42E-06 |
| Lifetime smoking-WHRadjBMI | 0,07 | 0,02 | 1,44E-04 | -0,07 | 0,02 | 2,01E-05 |
| Lifetime smoking-WC | 0,10 | 0,02 | 7,03E-06 | -0,12 | 0,03 | 2,19E-05 |
| Lifetime smoking - WCadjBMI | 0,15 | 0,01 | 5,86E-27 | -0,003 | 0,01 | 7,77E-01 |
| Lifetime smoking-HC | 0,10 | 0,02 | 6,40E-11 | -0,11 | 0,02 | 3,24E-06 |
| Lifetime smoking - HCadjBMI | 0,13 | 0,01 | 1,74E-28 | 0,06 | 0,03 | 6,13E-02 |
| Smoking initiation-WHR | 0,06 | 0,01 | 8,84E-07 | -0,06 | 0,02 | 6,03E-04 |
| Smoking initiation-WHRadjBMI | 0,05 | 0,01 | 7,43E-04 | -0,06 | 0,02 | 7,04E-05 |
| Smoking initiation-WC | 0,08 | 0,01 | 1,46E-08 | -0,12 | 0,02 | 1,63E-06 |
| Smoking initiation-WCadjBMI | 0,13 | 0,01 | 8,06E-36 | -0,01 | 0,01 | 1,97E-01 |
| Smoking initiation-HC | 0,05 | 0,03 | 8,35E-02 | -0,09 | 0,04 | 1,23E-02 |
| Smoking initiation-HCadjBMI | 0,10 | 0,01 | 3,95E-20 | 0,05 | 0,02 | 2,37E-02 |
| Pack-years-WHR | 0,12 | 0,03 | 5,82E-05 | -0,12 | 0,02 | 7,62E-08 |
| Pack-years-WHRadjBMI | 0,10 | 0,03 | 4,24E-03 | -0,09 | 0,02 | 8,05E-08 |
| Pack-years-WC | 0,11 | 0,05 | 0,01 | -0,12 | 0,04 | 6,82E-04 |
| Pack-years - WCadjBMI | 0,22 | 0,02 | 7,98E-22 | 0,004 | 0,01 | 7,02E-01 |
| Pack-years-HC | 0,12 | 0,05 | 0,01 | -0,12 | 0,04 | 5,12E-03 |
| Pack-years - HCadjBMI | 0,15 | 0,03 | 1,20E-05 | 0,03 | 0,05 | 4,91E-01 |
| Cigarettes per day (past/current) -WHRadjBMI | 0,00 | 0,03 | 0,93 | -0,18 | 0,04 | 1,05E-05 |
| Cigarettes per day (past/current) -WCadjBMI | 0,04 | 0,05 | 0,42 | -0,05 | 0,08 | 5,58E-01 |
| Cigarettes per day (current) -WHRadjBMI | 0,01 | 0,12 | 0,93 | 0,00 | 0,13 | 9,92E-01 |
| Cigarettes per day (current) -WCadjBMI | 0,01 | 0,03 | 0,72 | 0,34 | 0,02 | 4,35E-55 |
| LHC-MR, Latent Heritable Confounder Mendelian randomization; WHR, waist-to-hip ratio; WHRAdjBMI, waist-to-hip ratio adjusted for body mass index; WC, waist circunference; WCadjBMI, waist circunference adjusted for body mass index; HC, hip circumference; HCadjBMI, hip circumference adjusted for body mass index; SE, standard error. | | | | | | |

| **Supplementary Table 5: LHC-MR bidirectional causal effects between smoking and adiposity traits** | | | | | | |
| --- | --- | --- | --- | --- | --- | --- |
|  | **Smoking causal effect on adiposity** | | | **Adiposity causal effect on smoking** | | |
| **Exposure-outcome pairs** | **Coefficient** | **SE** | **P-value** | **Coefficient** | **SE** | **P-value** |
| Lifetime smoking-WHR | 0.44 | 0.03 | 5.84E-40 | 0.08 | 0.02 | 5.56E-04 |
| Lifetime smoking-WHRadjBMI | 0.26 | 0.03 | 1.65E-19 | -0.01 | 0.02 | 0.75 |
| Lifetime smoking-WC | 0.42 | 0.11 | 1.12E-04 | 0.14 | 0.06 | 0.01 |
| Lifetime smoking-WCadjBMI | 0.02 | 0.02 | 0.47 | 0.07 | 0.04 | 0.11 |
| Lifetime smoking-HC | 0.34 | 0.07 | 2.99E-07 | 0.10 | 0.03 | 4.60E-04 |
| Lifetime smoking-HCadjBMI | -0.17 | 0.03 | 3.23E-07 | -0.03 | 0.02 | 0.14 |
| Smoking initiation-WHR | 0.49 | 0.04 | 5.58E-33 | 0.01 | 0.02 | 0.63 |
| Smoking initiation-WHRadjBMI | 0.31 | 0.03 | 1.70E-29 | -0.02 | 0.01 | 0.13 |
| Smoking initiation-WC | 0.37 | 0.12 | 1.77E-03 | 0.12 | 0.04 | 3.33E-03 |
| Smoking initiation-WCadjBMI | 0.07 | 0.02 | 5.32E-04 | 0.03 | 0.03 | 0.38 |
| Smoking initiation-HC | 0.15 | 0.22 | 5.02E-01 | 0.11 | 0.04 | 9.21E-03 |
| Smoking initiation-HC | -0.08 | 0.03 | 9.05E-03 | 4.67E-03 | 0.02 | 0.76 |
| Pack-years-WHR | 0.37 | 0.10 | 3.26E-04 | 0.27 | 0.03 | 3.58E-18 |
| Pack-years-WHRadjBMI | 0.24 | 0.05 | 7.90E-06 | 0.07 | 0.02 | 2.53E-03 |
| Pack-years-WC | 0.25 | 0.20 | 0.22 | 0.32 | 0.04 | 5.14E-16 |
| Pack-years-WCadjBMI | -0.01 | 0.02 | 7.41E-01 | 0.16 | 0.05 | 4.50E-03 |
| Pack-years-HC | 0.43 | 0.20 | 0.04 | 0.17 | 0.05 | 1.85E-03 |
| Pack-years-HCadjBMI | -0.13 | 0.06 | 0.04 | -0.02 | 0.07 | 0.81 |
| Cigarettes per day (past/current)-WHRadjBMI | 0.06 | 0.10 | 0.55 | 0.24 | 0.07 | 2.75E-04 |
| Cigarettes per day (past/current)-WCadjBMI | -0.11 | 0.16 | 0.48 | 0.15 | 0.07 | 0.04 |
| Cigarettes per day (current)-WHRadjBMI | 0.24 | 0.16 | 0.13 | 0.25 | 0.28 | 0.37 |
| Cigarettes per day (current)-WCadjBMI | -0.36 | 0.19 | 0.06 | 0.33 | 0.11 | 3.34E-03 |
| LHC-MR, Latent Heritable Confounder Mendelian randomization; WHR, waist-hip ratio; WHRAdjBMI, waist-hip ratio adjusted for body mass index; WC, waist circumference; WCadjBMI, waist circumference adjusted for body mass index; HC, hip circumference; HCadjBMI, hip circumference adjusted for body mass index; SE, standard error. | | | | | | |

| **Supplementary table 6. Mendelian randomization results using the IVW, Egger, weighted median and weighted mode methods** | | | | | | | | | | | | | | | | | | | | | | | | | |
| --- | --- | --- | --- | --- | --- | --- | --- | --- | --- | --- | --- | --- | --- | --- | --- | --- | --- | --- | --- | --- | --- | --- | --- | --- | --- |
|  |  | **WHR** | | | | **WHRAdjBMI** | | | | **WC** | | | | **WCAdjBMI** | | | | **HC** | | | | **HCadjBMI** | | | |
| **Exposure** | **Model** | **SNP** | **Beta** | **SE** | **p-value** | **SNP** | **beta** | **SE** | **p-value** | **SNP** | **beta** | **SE** | **p-value** | **SNP** | **beta** | **SE** | **p-value** | **SNP** | **beta** | **SE** | **p-value** | **SNP** | **beta** | **SE** | **p-value** |
| **Lifetime smoking** | IVW | 83 | 0.18 | 0.05 | 2.28E-04 | 83 | 0.18 | 0.05 | 0.0004 | 80 | 0.06 | 0.06 | 0.33 | 86 | 0.14 | 0.05 | 2.74E-03 | 81 | 0.01 | 0.05 | 0.87 | 89 | -0.05 | 0.05 | 0.36 |
|  | Egger | 83 | 0.06 | 0.21 | 0.78 | 83 | 0.32 | 0.20 | 0.12 | 80 | -0.18 | 0.22 | 0.40 | 86 | 0.33 | 0.19 | 0.09 | 81 | -0.30 | 0.21 | 0.16 | 89 | 0.09 | 0.21 | 0.66 |
|  | Weighted median | 83 | 0.20 | 0.08 | 1.00E-02 | 83 | 0.12 | 0.08 | 0.11 | 80 | 0.02 | 0.08 | 0.80 | 86 | 0.17 | 0.07 | 0.02 | 81 | -0.04 | 0.08 | 0.59 | 89 | -0.04 | 0.08 | 0.65 |
|  | Weighted mode | 83 | 0.17 | 0.21 | 0.44 | 83 | -0.06 | 0.16 | 0.72 | 80 | -0.12 | 0.17 | 0.47 | 86 | 0.23 | 0.13 | 0.08 | 81 | -0.28 | 0.18 | 0.12 | 89 | -0.23 | 0.27 | 0.39 |
| **Smoking initiation** | IVW | 121 | 0.07 | 0.02 | 8.96E-04 | 130 | 0.03 | 0.02 | 0.15 | 115 | 0.06 | 0.02 | 0.01 | 131 | 3.51E-03 | 0.02 | 0.87 | 116 | 0.02 | 0.02 | 0.48 | 127 | -0.01 | 0.02 | 0.77 |
|  | Egger | 121 | 0.19 | 0.10 | 0.06 | 130 | 0.13 | 0.09 | 0.16 | 115 | 0.07 | 0.10 | 0.52 | 131 | 0.15 | 0.09 | 0.10 | 116 | -0.01 | 0.11 | 0.90 | 127 | 0.03 | 0.10 | 0.78 |
|  | Weighted median | 121 | 0.12 | 0.03 | 1.15E-04 | 130 | 0.03 | 0.03 | 0.29 | 115 | 0.09 | 0.03 | 0.01 | 131 | -0.02 | 0.03 | 0.61 | 116 | 0.03 | 0.03 | 0.35 | 127 | -0.03 | 0.03 | 0.41 |
|  | Weighted mode | 121 | 0.19 | 0.09 | 0.03 | 130 | -0.02 | 0.08 | 0.85 | 115 | 0.11 | 0.09 | 0.21 | 131 | 0.19 | 0.10 | 0.05 | 116 | 0.05 | 0.09 | 0.57 | 127 | -0.07 | 0.10 | 0.50 |
| **Pack years of adult smoking** | IVW | 9 | 0.03 | 0.05 | 0.55 | 10 | 0.08 | 0.04 | 0.04 | 8 | -0.22 | 0.10 | 0.07 | 10 | 0.03 | 0.05 | 0.58 | 8 | -0.06 | 0.06 | 0.34 | 10 | -0.05 | 0.05 | 0.33 |
|  | Egger | 9 | -0.10 | 0.08 | 0.26 | 10 | 0.01 | 0.08 | 0.93 | 8 | -0.22 | 0.10 | 0.07 | 10 | 0.02 | 0.11 | 0.89 | 8 | -0.21 | 0.11 | 0.10 | 10 | 0.03 | 0.10 | 0.76 |
|  | Weighted median | 9 | -2.44E-03 | 0.04 | 0.95 | 10 | 0.05 | 0.04 | 0.20 | 8 | -0.06 | 0.04 | 0.16 | 10 | 0.02 | 0.05 | 0.71 | 8 | -0.11 | 0.05 | 0.03 | 10 | -0.05 | 0.05 | 0.32 |
|  | Weighted mode | 9 | -0.02 | 0.04 | 0.62 | 10 | 0.05 | 0.05 | 0.34 | 8 | -0.08 | 0.05 | 0.13 | 10 | 0.01 | 0.05 | 0.81 | 8 | -0.11 | 0.05 | 0.05 | 10 | -0.05 | 0.05 | 0.34 |
| **Cigarettes per day (past/current)** | IVW | NA | NA | NA | NA | 12 | 0.03 | 0.04 | 0.50 | NA | NA | NA | NA | 13 | 0.10 | 0.07 | 0.14 | NA | NA | NA | NA | NA | NA | NA | NA |
|  | Egger | NA | NA | NA | NA | 12 | -0.01 | 0.07 | 0.87 | NA | NA | NA | NA | 13 | 0.12 | 0.10 | 0.26 | NA | NA | NA | NA | NA | NA | NA | NA |
|  | Weighted median | NA | NA | NA | NA | 12 | 0.02 | 0.04 | 0.64 | NA | NA | NA | NA | 13 | 0.11 | 0.08 | 0.15 | NA | NA | NA | NA | NA | NA | NA | NA |
|  | Weighted mode | NA | NA | NA | NA | 12 | 0.02 | 0.05 | 0.72 | NA | NA | NA | NA | 13 | 0.11 | 0.08 | 0.17 | NA | NA | NA | NA | NA | NA | NA | NA |
| **Cigarettes per day (current)** | IVW | NA | NA | NA | NA | 3 | 0.24 | 0.10 | 0.02 | NA | NA | NA | NA | 3 | 0.11 | 0.18 | 0.53 | NA | NA | NA | NA | NA | NA | NA | NA |
|  | Egger | NA | NA | NA | NA | 3 | 0.22 | 0.29 | 0.59 | NA | NA | NA | NA | 3 | 0.60 | 0.50 | 0.44 | NA | NA | NA | NA | NA | NA | NA | NA |
|  | Weighted median | NA | NA | NA | NA | 3 | 0.24 | 0.10 | 0.02 | NA | NA | NA | NA | 3 | 0.20 | 0.11 | 0.07 | NA | NA | NA | NA | NA | NA | NA | NA |
|  | Weighted mode | NA | NA | NA | NA | 3 | 0.23 | 0.12 | 0.18 | NA | NA | NA | NA | 3 | 0.21 | 0.11 | 0.20 | NA | NA | NA | NA | NA | NA | NA | NA |
| WHR, waist-hip ratio; WHRAdjBMI, waist-hip ratio adjusted for body mass index; WC, waist circumference; WCadjBMI, waist circumference adjusted for body mass index; HC, hip circumference; HCadjBMI, hip circumference adjusted for body mass index; SE, standard error; NA, not available. | | | | | | | | | | | | | | | | | | | | | | | | | |

| S**upplementary Table 7. Sensitivity tests for IVW, Egger, weighted median and weighted mode Mendelian randomization methods** | | | | | | | | | | | | | |
| --- | --- | --- | --- | --- | --- | --- | --- | --- | --- | --- | --- | --- | --- |
|  |  | **WHR** | | **WHRAdjBMI** | | **WC** | | **WCAdjBMI** | | **HC** | | **HCadjBMI** | |
| **Smoking trait** | **Sensitivity test** | **Estimate** | **p-value/CI** | **Estimate** | **p-value/CI** | **Estimate** | **p-value/CI** | **Estimate** | **p-value/CI** | **Estimate** | **p-value/CI** | **Estimate** | **p-value/CI** |
| Lifetime smoking | Q (p-value) | 87.29 | 0.32 | 80.69 | 0.49 | 90.85 | 0.17 | 73.48 | 0.81 | 73.23 | 0.66 | 94.68 | 0.27 |
|  | I2 (CI) | 6.10 | 0%; 28,60% | 0 | 0%; 26,40% | 13.00 | 0%; 34,80% | 0 | 0%; 14,50% | 0 | 0%; 21,20% | 8,10% | 0%; 30,10% |
|  | Q-Q' (p-value) | 0.47 | 0.50 | 0.57 | 0.45 | 1.57 | 0.21 | 0.97 | 0.32 | 2.29 | 0.13 | 0.57 | 0.47 |
|  | Egger intercept | 1,36-03 | 0.50 | -1.56E-03 | 0.45 | 2.52E-03 | 0.21 | -1.98E-03 | 0.30 | 3.29E-03 | 0.12 | -1.51E-03 | 0.47 |
|  | Rucker test | IVW | | IVW | | IVW | | IVW | | IVW | | IVW | |
| Smoking initiation | Q (p-value) | 70.77 | 1.00 | 70.11 | 1 | 65.16 | 1.00 | 87.20 | 1.00 | 68.44 | 1.00 | 80.59 | 0.9994 |
|  | I2 (CI) | 0 | 0%; 0% | 0 | 0%; 0% | 0 | 0%; 0% | 0 | 0%; 0% | 0 | 0%; 0% | 0 | 0%; 0% |
|  | Q-Q' (p-value) | 1.45 | 0.23 | 1.26 | 0.26 | 0.01 | 0.91 | 2.76 | 0.10 | 0.08 | 0.78 | 0.13 | 0.72 |
|  | Egger intercept | -2.22E-03 | 0.12 | -1.94E-03 | 0.13 | -2.18E-04 | 0.88 | -2.85E-03 | 0.04 | 5.68E-04 | 0.72 | -6.55E-04 | 0.66 |
|  | Rucker test | IVW | | IVW | | IVW | | IVW | | IVW | | IVW | |
| Pack-years | Q (p-value) | 11.56 | 0.1717 | 8.82 | 0.4540 | 15.20 | 0.03 | 16.91 | 0.05 | 13.69 | 0.06 | 12.15% | 0.2047 |
|  | I2 (CI) | 30.80 | 0%; 68,00% | 0 | 0%; 62.40% | 53.90 | 0; 79,20% | 46.80% | 0%; 74,40% | 48,80% | 0%; 77,20% | 26.00% | 0,00%; 64,30% |
|  | Q-Q' (p-value) | 3.94 | 4.72E-02 | 1.33 | 0.25 | 0.01 | 5.35E-03 | 0.05 | 0.83 | 4.41 | 0.04 | -3.51E-03 | 0.25 |
|  | Egger intercept | 6.25E-03 | 4.72E-02 | 3.37E-03 | 0.25 | 7.76 | 5.35E-03 | 6.35E-04 | 0.83 | 7.81E-03 | 0.04 | 1.31 | 0.25 |
|  | Rucker test | IVW | | IVW | | IVW | | IVW | | IVW | | IVW | |
| Cigarettes per day (past/current) | Q (p-value) | NA | NA | 6.82 | 0.81 | NA | NA | 9.35 | 0.67 | NA | NA | NA | NA |
|  | I2 (CI) | NA | NA | 0 | 0,00%; 32,80% | NA | NA | 0% | 0,00%; 44,30% | NA | NA | NA | NA |
|  | Q-Q' (p-value) | NA | NA | 0.58 | 0.45 | NA | NA | 0.06 | 0.81 | NA | NA | NA | NA |
|  | Egger intercept | NA | NA | 2.95E-03 | 0.34 | NA | NA | -9.32E-04 | 0.80 | NA | NA | NA | NA |
|  | Rucker test | NA | | IVW | | NA | | IVW | | NA | | NA | |
| Cigarettes per day (current) | Q (p-value) | NA | NA | 0.17 | 0.92 | NA | NA | 5.76 | 0.06 | NA | NA | NA | NA |
|  | I2 (CI) | NA | NA | 0 | 0%; 0% | NA | NA | 65,30% | 0,00%; 90,00% | NA | NA | NA | NA |
|  | Q-Q' (p-value) | NA | NA | 3.91E-03 | 0.95 | NA | NA | 3.42 | 0.06 | NA | NA | NA | NA |
|  | Egger intercept | NA | NA | 1.04E-03 | 0.88 | NA | NA | -0.03 | 0.06 | NA | NA | NA | NA |
|  | Rucker test | NA | | IVW | | NA | | IVW | | NA | | NA | |
| WHR, waist-hip ratio; WHRAdjBMI, waist- hip ratio adjusted for body mass index; WC, waist circumference; WCadjBMI, waist circumference adjusted for body mass index; HC, hip circumference; HCadjBMI, hip circumference adjusted for body mass index; NA, not available. | | | | | | | | | | | | | |

| **Supplementary Table 8: Multivariable Mendelian randomization results between lifetime smoking and education with abdominal adiposity traits** | | | | | | |
| --- | --- | --- | --- | --- | --- | --- |
| **Trait** | **SNP** | **mF** | **Cochran's Q (P-value)** | **beta** | **SE** | **P-value** |
| **WHR** | | | | | | |
| Lifetime Smoking | 207 | 11.04 | 163.70 | 0.36 | 0.06 | 9,43E-10 |
| Education | 207 | 18.87 | 0.98 | -0.28 | 0.05 | 2,09E-07 |
| **WHRadjBMI** | | | | | | |
| Lifetime Smoking | 212 | 12.02 | 190.54 | 0.29 | 0.06 | 6,27E-07 |
| Education | 212 | 19.64 | 0.82 | -0.12 | 0.06 | 0.03 |
| **WC** | | | | | | |
| Lifetime Smoking | 199 | 10.27 | 184.07 | 0.38 | 0.07 | 2.68E-08 |
| Education | 199 | 17.54 | 0.72 | -0.27 | 0.06 | 1.19E-05 |
| **WCadjBMI** | | | | | | |
| Lifetime Smoking | 204 | 12.55 | 169.82 | 0.22 | 0.05 | 5.91E-05 |
| Education | 204 | 20.89 | 0.95 | 0.09 | 0.05 | 0.11 |
| **HC** | | | | | | |
| Lifetime Smoking | 195 | 10.16 | 181.14 | 0.19 | 0.07 | 6.36E-03 |
| Education | 195 | 17.58 | 0.70 | -0.15 | 0.06 | 0.02 |
| **HCadjBMI** | | | | | | |
| Lifetime Smoking | 199 | 12.80 | 183.57 | -0.05 | 0.06 | 0.36 |
| Education | 199 | 21.77 | 0.73 | 0.20 | 0.06 | 8.60E-04 |
| WHR, waist-to-hip ratio; WHRAdjBMI, waist-to-hip ratio adjusted for body mass index; WC, waist circumference; WCadjBMI, waist circumference adjusted for body mass index; HC, hip circumference; HCadjBMI, hip circumference adjusted for body mass index. | | | | | | |

| **Supplementary Table 9: Multivariable Mendelian randomization results between smoking initiation and education with abdominal adiposity traits** | | | | | | |
| --- | --- | --- | --- | --- | --- | --- |
| **Trait** | **SNP** | **mF** | **Cochran's Q (P-value)** | **beta** | **SE** | **P-value** |
| **WHR** | | | | | | |
| Smoking initiation | 207 | 10.57 | 167.30 | 0.10 | 0.02 | 1.28E-05 |
| Education | 207 | 28.25 | 0.97 | -0.38 | 0.05 | 9.74E-14 |
| **WHRadjBMI** | | | | | | |
| Smoking initiation | 207 | 11.33 | 164.67 | 0.09 | 0.02 | 1.30E-04 |
| Education | 207 | 27.34 | 0.98 | -0.15 | 0.05 | 2.70E-03 |
| **WC** | | | | | | |
| Smoking initiation | 195 | 10.19 | 179.54 | 0.08 | 0.03 | 2.31E-03 |
| Education | 195 | 26.39 | 0.73 | -0.36 | 0.05 | 3.29E-10 |
| **WCadjBMI** | | | | | | |
| Smoking initiation | 202 | 10.84 | 167.31 | 0.06 | 0.02 | 7.59E-03 |
| Education | 202 | 26.64 | 0.95 | 0.05 | 0.05 | 2.71E-01 |
| **HC** | | | | | | |
| Smoking initiation | 194 | 10.56 | 189.92 | 0.05 | 0.03 | 7.41E-02 |
| Education | 194 | 27.84 | 0.51 | -0.27 | 0.06 | 5.84E-06 |
| **HCadjBMI** | | | | | | |
| Smoking initiation | 196 | 11.27 | 164.15 | -0.01 | 0.02 | 0.63 |
| Education | 196 | 27.71 | 0.94 | 0.19 | 0.05 | 3.57E-04 |
| WHR, waist-to-hip ratio; WHRAdjBMI, waist-to-hip ratio adjusted for body mass index; WC, waist circumference; WCadjBMI, waist circumference adjusted for body mass index; HC, hip circumference; HCadjBMI, hip circumference adjusted for body mass index. | | | | | | |

| **Supplementary Table 10: Wald ratio estimates for the association between smoking and body fat distribution traits for the rs1317286 variant in the *CHRNA5/3* locus** | | | | | | | | | | | | | | | | | | | |
| --- | --- | --- | --- | --- | --- | --- | --- | --- | --- | --- | --- | --- | --- | --- | --- | --- | --- | --- | --- |
|  |  | **WHR** | | | **WHRadjBMI** | | | **WC** | | | **WCadjBMI** | | | **HC** | | | **HCadjBMI** | | |
| **Smoking trait** | **F-statistic** | **beta** | **SE** | **p-value** | **beta** | **SE** | **p-value** | **beta** | **SE** | **p-value** | **beta** | **SE** | **p-value** | **beta** | **SE** | **p-value** | **beta** | **SE** | **p-value** |
| Lifetime smoking | 157.94 | 0.13 | 0.24 | 0.59 | 0.49 | 0.24 | 0.04 | -0.29 | 0.24 | 0.24 | 0.32 | 0.24 | 0.17 | -0.50 | 0.25 | 0.05 | -0.16 | 0.25 | 0.51 |
| Smoking initiation | 6.78 | -0.24 | 0.45 | 0.59 | -0.92 | 0.45 | 0.04 | 0.54 | 0.46 | 0.24 | -0.61 | 0.45 | 0.17 | 0.94 | 0.47 | 0.05 | 0.31 | 0.47 | 0.51 |
| Pack years adult smoking | 401.53 | 0.03 | 0.06 | 0.59 | 0.12 | 0.06 | 0.04 | -0.07 | 0.06 | 0.24 | 0.08 | 0.06 | 0.17 | -0.13 | 0.06 | 0.05 | -0.04 | 0.06 | 0.51 |
| Cigarettes per day (past/current) | 925.99 | NA | NA | NA | 0.10 | 0.05 | 0.04 | NA | NA | NA | 0.09 | 0.05 | 0.06 | NA | NA | NA | NA | NA | NA |
| Cigarettes per day (current) | 178.58 | NA | NA | NA | 0.24 | 0.12 | 0.04 | NA | NA | NA | 0.22 | 0.11 | 0.06 | NA | NA | NA | NA | NA | NA |
| WHR, waist-to-hip ratio; WHRAdjBMI, waist-to-hip ratio adjusted for body mass index; WC, waist circumference; WCadjBMI, waist circumference adjusted for body mass index; HC, hip circumference; HCadjBMI, hip circumference adjusted for body mass index; SE, standard error; NA, not available. | | | | | | | | | | | | | | | | | | | |

| **Supplementary Table 11: Association of genetic risk scores for smoking traits with fat depot volumes and their ratios.** | | | | | | | | | | | | | | |
| --- | --- | --- | --- | --- | --- | --- | --- | --- | --- | --- | --- | --- | --- | --- |
|  | **Lifetime smoking - WHR** | | | | | |  | | **Lifetime smoking - WHRadjBMI** | | | | | |
| **Fat depot** | **N** | **effect size** | **SE** | **lower 95% CI** | **upper 95% CI** | **P** | **Fat depot** | **N** | | **effect size** | **SE** | **lower 95%**  **CI** | **upper**  **95% CI** | **P** |
| VAT | 76 | 0.31 | 0.09 | 0.13 | 0.48 | 5.08E-04 | VATadjBMI | 83 | | 0.1 | 0.08 | -0.06 | 0.27 | 2.08E-01 |
| GSAT | 76 | 0.26 | 0.09 | 0.09 | 0.43 | 2.97E-03 | GSATadjBMI | 83 | | -0.003 | 0.08 | -0.16 | 0.16 | 9.75E-01 |
| ASAT | 76 | 0.21 | 0.09 | 0.04 | 0.39 | 1.56E-02 | ASATadjBMI | 83 | | -0.13 | 0.08 | -0.30 | 0.03 | 1.15E-01 |
| VAT/GSAT | 76 | 0.22 | 0.09 | 0.05 | 0.39 | 1.00E-02 | VAT/GSAT | NA | | NA | NA | NA | NA | NA |
| VAT/ASAT | 76 | 0.25 | 0.09 | 0.08 | 0.42 | 3.17E-03 | VAT/ASAT | NA | | NA | NA | NA | NA | NA |
| ASAT/GSAT | 76 | 0.06 | 0.09 | -0.11 | 0.23 | 4.78E-01 | ASAT/GSAT | NA | | NA | NA | NA | NA | NA |
|  | **Smoking initiation - WHR** | | | | | |  | | **Smoking initiation - WHRadjBMI** | | | | | |
| VAT | 121 | 0.13 | 0.04 | 0.06 | 0.20 | 5.66E-04 | VATadjBMI | 130 | | -0.04 | 0.04 | -0.11 | 0.03 | 2.89E-01 |
| GSAT | 121 | 0.10 | 0.04 | 0.02 | 0.17 | 8.98E-03 | GSATadjBMI | 130 | | -0.03 | 0.04 | -0.10 | 0.04 | 3.50E-01 |
| ASAT | 121 | 0.09 | 0.04 | 0.02 | 0.16 | 1.65E-02 | ASATadjBMI | 130 | | -0.11 | 0.04 | -0.18 | -0.04 | 3.54E-03 |
| VAT/GSAT | 121 | 0.10 | 0.04 | 0.03 | 0.17 | 5.93E-03 | VAT/GSAT | NA | | NA | NA | NA | NA | NA |
| VAT/ASAT | 121 | 0.10 | 0.04 | 0.02 | 0.17 | 8.42E-03 | VAT/ASAT | NA | | NA | NA | NA | NA | NA |
| ASAT/GSAT | 121 | 0.05 | 0.04 | -0.03 | 0.12 | 2.12E-01 | ASAT/GSAT | NA | | NA | NA | NA | NA | NA |
| We constructed genetic risk scores using variants from the inverse variance-weighted model for the causal association between smoking initiation and WHR (121 variants), smoking initiation and WHR_adjBMI_ (130 variants), lifetime smoking and WHR (76 variants), and lifetime smoking and WHR_adjBMI_ (83 variants). The smoking variants causally associated with WHR were used to study associations with BMI-unadjusted fat depots, while the smoking variants causally associated with WHR_adjBMI_ were used to study associations with BMI-adjusted fat depots. We computed weighted genetic scores using the beta of the variants for a smoking trait as weights with gtx.package’s (v0.0.8) grs.summary function. This function approximates the results of a genetic risk score using GWAS summary statistics by calculating the joint effect of genetic variants on an outcome trait. N, number of variants used; SE, standard error; P, p-value; CI, confidence interval; VAT, visceral adipose tissue; ASAT, abdominal subcutaneous adipose tissue; GSAT, gluteofemoral subcutaneous adipose tissue. | | | | | | | | | | | | | | |

| **Supplementary Table 12: Association of genetic risk scores for smoking traits with hormones.** | | | | | | |
| --- | --- | --- | --- | --- | --- | --- |
|  | **Lifetime smoking - WHRadjBMI** | | | | | |
| **Hormones** | **N** | **effect size** | **SE** | **lower  95% CI** | **upper**  **95% CI** | **P** |
| Cortisol | 69 | -0.01 | 0.11 | -0.23 | 0.22 | 9.60E-01 |
| Oestradiol | 69 | 0.12 | 0.11 | -0.10 | 0.33 | 2.90E-01 |
| Testosterone | 69 | -0.01 | 0.02 | -0.04 | 0.03 | 7.12E-01 |
| SHBG | 69 | -0.002 | 0.01 | -0.02 | 0.02 | 8.79E-01 |
|  | **Smoking initiation - WHRadjBMI** | | | | | |
| Cortisol | 110 | -0.07 | 0.05 | -0.17 | 0.03 | 1.77E-01 |
| Oestradiol | 110 | 0.01 | 0.05 | -0.08 | 0.11 | 7.74E-01 |
| Testosterone | 110 | -0.01 | 0.01 | -0.02 | 0.01 | 3.02E-01 |
| SHBG | 110 | -0.02 | 0.01 | -0.03 | -0.01 | 4.56E-05 |
| We constructed genetic risk scores using variants from the inverse variance-weighted model for the causal association between smoking initiation and WHR_adjBMI_ (110 variants), and lifetime smoking and WHR_adjBMI_ (69 variants). We computed weighted genetic scores using the beta of the variants for a smoking trait as weights with gtx.package’s (v0.0.8) grs.summary function. This function approximates the results of a genetic risk score using GWAS summary statistics by calculating the joint effect of genetic variants on an outcome trait. N, number of variants used; SE, standard error; P, p-value; CI, confidence interval. | | | | | | |

**STROBE-MR checklist of recommended items to address in reports of Mendelian randomization studies**^1^ ^2^

| **Item No.** | **Section** | **Checklist item** | **Page No.** | **Relevant text from manuscript** |
| --- | --- | --- | --- | --- |
| 1 | **TITLE and ABSTRACT** | Indicate Mendelian randomization (MR) as the study’s design in the title and/or the abstract if that is a main purpose of the study | 1  2 | TITLE “Estimating causality between smoking and abdominal obesity by Mendelian randomization”  ABSTRACT “We implemented Mendelian randomization using the CAUSE and LHC-MR methods that instrument smoking using genome-wide data” |
|  | **INTRODUCTION** |  |  |  |
| 2 | **Background** | Explain the scientific background and rationale for the reported study. What is the exposure? Is a potential causal relationship between exposure and outcome plausible? Justify why MR is a helpful method to address the study question | 3 | “Whether the association between smoking and body fat distribution is causal or explained by confounding or reverse causality is still unclear.” “Studies based on a single genetic instrument are sensitive to genetic pleiotropy and may produce imprecise causal estimates, leaving the relationship between smoking and abdominal obesity uncertain.” |
| 3 | **Objectives** | State specific objectives clearly, including pre-specified causal hypotheses (if any). State that MR is a method that, under specific assumptions, intends to estimate causal effects | 3 | “Here, we report on two-sample Mendelian randomization analyses to estimate the causal effect of smoking initiation, smoking heaviness, and lifetime smoking (which captures smoking heaviness, duration, and time of cessation) on abdominal adiposity.” |
|  | **METHODS** |  |  |  |
| 4 | **Study design and data sources** | Present key elements of the study design early in the article. Consider including a table listing sources of data for all phases of the study. For each data source contributing to the analysis, describe the following: | 3-5 | Section “Mendelian randomization methods” |
|  | a) | Setting: Describe the study design and the underlying population, if possible. Describe the setting, locations, and relevant dates, including periods of recruitment, exposure, follow-up, and data collection, when available. | 5-6 | “Data sources for smoking and body fat distribution” |
|  | b) | Participants: Give the eligibility criteria, and the sources and methods of selection of participants. Report the sample size, and whether any power or sample size calculations were carried out prior to the main analysis | 5 | “We utilized the largest published GWAS summary level data of European ancestry”  “We also utilized the largest published GWAS summary level data of European ancestry for measures of body fat distribution” |
|  | c) | Describe measurement, quality control and selection of genetic variants | 20-21 | Section “Mendelian randomization using the CAUSE Method” Section “Mendelian randomization using the LHC-MR method”  Section “Mendelian randomization using the inverse variance-weighted (IVW), MR-Egger, weighted median, and weighted mode methods” |
|  | d) | For each exposure, outcome, and other relevant variables, describe methods of assessment and diagnostic criteria for diseases | 5-6 | Section “Data sources for smoking and body fat distribution” Section “Association of smoking variants with body fat depot volumes and hormonal levels” |
|  | e) | Provide details of ethics committee approval and participant informed consent, if relevant | 6 | “The present study used public GWAS summary-level data and did not have direct contact with study participants. Therefore, no direct ethical approval was needed.” |
| 5 | **Assumptions** | Explicitly state the three core IV assumptions for the main analysis (relevance, independence and exclusion restriction) as well assumptions for any additional or sensitivity analysis | 12 | Figure 1 and Figure 1 legend |
| 6 | **Statistical methods: main analysis** | Describe statistical methods and statistics used | 20-21 | Section “Mendelian randomization using the CAUSE Method”  Section “Mendelian randomization using the LHC-MR method”  Section “Mendelian randomization using the inverse variance-weighted (IVW), MR-Egger, weighted median, and weighted mode methods” |
|  | a) | Describe how quantitative variables were handled in the analyses (i.e., scale, units, model) | 5 | “… smoking initiation (n=1,232,091), defined as whether an individual has ever smoked regularly (yes/no) [17]; lifetime smoking (n=462,690), which captures the initiation, duration, heaviness, and time since the cessation of smoking [31]; and smoking heaviness, defined as cigarettes smoked per day by current smokers only or current and former smokers combined (n=337,334) [17], or as the total number of pack-years smoked in adulthood (n = 142,387)”  “WHR (n=697,734) and WHR adjusted for body mass index (WHRadjBMI, n=694,649) from a meta-analysis of the Genetic Investigation of Anthropometric Traits (GIANT) consortium and the UK Biobank [32]; waist and hip circumferences (WC and HC) from the UK Biobank (n=462,166 and n=462,177, respectively); and WCadjBMI and HCadjBMI from the GIANT consortium (n=231,355 and n=211,117, respectively)” |
|  | b) | Describe how genetic variants were handled in the analyses and, if applicable, how their weights were selected | 20-21 | Section “Mendelian randomization using the CAUSE Method”  Section “Mendelian randomization using the LHC-MR method”  Section “Mendelian randomization using the inverse variance-weighted (IVW), MR-Egger, weighted median, and weighted mode methods”” |
|  | c) | Describe the MR estimator (e.g. two-stage least squares, Wald ratio) and related statistics. Detail the included covariates and, in case of two-sample MR, whether the same covariate set was used for adjustment in the two samples | 3-5 | Section “Mendelian randomization methods” |
|  | d) | Explain how missing data were addressed | 23 | “We replaced the independent lead variants not present in the outcome trait GWAS summary statistics by proxy variants (r2 > 0.8 in European 1000 Genomes reference panel). To identify the most accurate proxy, we clumped (clump_kb = 10000kb and clump_r2 < 0.001) all proxies found in the locus of the missing lead variant, using the p-value for the exposure trait for which the missing lead variant was genome-wide significant. If the missing lead variant was found to be genome-wide significant for more than one trait, the p-value of the exposure trait for which the missing variant presented the lowest p-value was used.” |
|  | e) | If applicable, indicate how multiple testing was addressed |  | Not applicable |
| 7 | **Assessment of assumptions** | Describe any methods or prior knowledge used to assess the assumptions or justify their validity | 4-5 | “…applied Steiger filtering to remove variants that are likely to affect the outcome trait through other traits than the exposure trait. The strength of the genetic instruments was assessed using the F statistic…  We estimated heterogeneity across the causal estimates of SNPs using the Meta R package [23]. The causal estimates were considered heterogeneous if the P value for Cochran’s Q test was significant (P<0.05) and I2 was above 25%. We assessed bias introduced by horizontal pleiotropy by implementing the Egger’s intercept test. Egger’s intercept P<0.05 was considered as evidence of horizontal pleiotropy. We used the Rucker framework [24] to assess whether MR-Egger regression, which accounts for horizontal pleiotropy but limits statistical power, should be applied instead of the standard inverse variance-weighted model. We visually assessed heterogeneity and horizontal pleiotropy using leave-one-out forest plots and funnel plots. To detect individual pleiotropic variants that might bias the results, we used the RadialMR package in R (1.0.0) [25], applying an iterative Cochran’s Q method and setting a strict outlier threshold of P<0.05. The iterative Cochran’s Q, either inverse variance-weighted Q or Egger Q, was chosen depending on the Rucker framework results. After removing outlier variants detected with RadialMR, we re-ran the Mendelian randomization and sensitivity tests to ensure that horizontal pleiotropy introduced by the outlier variants had been removed.” |
| 8 | **Sensitivity analyses and additional analyses** | Describe any sensitivity analyses or additional analyses performed (e.g. comparison of effect estimates from different approaches, independent replication, bias analytic techniques, validation of instruments, simulations) | 20 | “For the inverse variance-weighted, MR-Egger, weighted median, and weighted mode Mendelian randomization methods, we applied a range of sensitivity tests (Supplementary Table 6) and plots (Supplementary Figure 3) to assess pleiotropy and ensure that the reported causal effects were unbiased.” |
| 9 | **Software and pre-registration** |  |  |  |
|  | a) | Name statistical software and package(s), including version and settings used | 4 | “In addition to the CAUSE (v1.0.0) and LHC-MR (v.1.0.0) methods … The analyses were performed using the TwoSampleMR (v.0.5.6) package in R [18]” |
|  | b) | State whether the study protocol and details were pre-registered (as well as when and where) |  | Not preregistered |
|  | **RESULTS** |  |  |  |
| 10 | **Descriptive data** |  |  |  |
|  | a) | Report the numbers of individuals at each stage of included studies and reasons for exclusion. Consider use of a flow diagram | 31 | “Supplementary Table 1: Datasets for smoking and body fat distribution traits used in the present study” |
|  | b) | Report summary statistics for phenotypic exposure(s), outcome(s), and other relevant variables (e.g. means, SDs, proportions) | 31 | “Supplementary Table 1: Datasets for smoking and body fat distribution traits used in the present study” |
|  | c) | If the data sources include meta-analyses of previous studies, provide the assessments of heterogeneity across these studies |  | Not applicable |
|  | d) | For two-sample MR:  i.  Provide justification of the similarity of the genetic variant-exposure associations between the exposure and outcome samples  ii.  Provide information on the number of individuals who overlap between the exposure and outcome studies | 5-6 | “The Causal Analysis Using the Summary Effect estimates (CAUSE) and the Latent Heritable Confounder MR (LHC-MR) methods utilize genome-wide association data to assess causal relationships rather than genome-wide significant loci only, to correct for sample overlap”  “In the Mendelian randomization analyses using the IVW, MR Egger, weighted median and weighted mode methods, which are sensitive to bias when there is a sample overlap between the exposure and outcome traits, we used the GIANT consortium data without the UK Biobank for each of the body fat distribution traits” |
| 11 | **Main results** |  |  |  |
|  | a) | Report the associations between genetic variant and exposure, and between genetic variant and outcome, preferably on an interpretable scale | 6-8 | Section “Effects of smoking initiation and lifetime smoking on abdominal adiposity”  “In original units, our results suggest that starting to smoke increases WHR by an average of 0.063 (95% CI: 0.012-0.114). This corresponds to an increase of approximately 7% in WHR, given that the average WHR in the population is 0.90.”  Section “Effect of smoking heaviness on abdominal adiposity in past and current smokers”  Section “Causal effect of abdominal obesity on smoking traits” |
|  | b) | Report MR estimates of the relationship between exposure and outcome, and the measures of uncertainty from the MR analysis, on an interpretable scale, such as odds ratio or relative risk per SD difference | 6-8 | Section “Effects of smoking initiation and lifetime smoking on abdominal adiposity”  In original units, our results suggest that starting to smoke increases WHR by an average of 0.063 (95% CI: 0.012-0.114). This corresponds to an increase of approximately 7% in WHR, given that the average WHR in the population is 0.90. “”  Section “Effect of smoking heaviness on abdominal adiposity in past and current smokers”  Section “Causal effect of abdominal obesity on smoking traits” |
|  | c) | If relevant, consider translating estimates of relative risk into absolute risk for a meaningful time period |  | Not relevant |
|  | d) | Consider plots to visualize results (e.g. forest plot, scatterplot of associations between genetic variants and outcome versus between genetic variants and exposure) | 4  21  25-27 | “We visually assessed heterogeneity and horizontal pleiotropy using leave-one-out forest plots and funnel plots.”  “We visualized the effects of heterogeneity and horizontal pleiotropy using leave-one-out forest plots and funnel plots (Supplementary Figure 3).  …  After the extraction of the outliers, the sensitivity tests and sensitivity plots showed no evidence of pleiotropy (Supplementary Table 6, Supplementary Figure 2 and 3)”  Supplementary figure 1,2 and 3 |
| 12 | **Assessment of assumptions** |  |  |  |
|  | a) | Report the assessment of the validity of the assumptions | 7 | “The CAUSE method showed no evidence of correlated horizontal pleiotropy, as indicated by the zero median shared effect and low q-values in all models (Supplementary Table 2). The LHC-MR results indicated an effect of confounders on WHR and WHRadjBMI, independent of smoking traits, suggesting the presence of correlated horizontal pleiotropy (Supplementary Tables 4-5). However, the reported confounder effect was directionally opposite to the positive causal effects of smoking initiation and lifetime smoking on WHR and WHRadjBMI, suggesting that the confounder effect leads to an underestimation of the true causal effects.”  Section “Interpretation of the results indicative of causality for inversed variant weighted (IVW), MR-Egger, weighted median, and weighted mode methods:” |
|  | b) | Report any additional statistics (e.g., assessments of heterogeneity across genetic variants, such as *I^2^*, Q statistic or E-value) | 21-23 | Section “Interpretation of the results indicative of causality for inversed variant weighted (IVW), MR-Egger, weighted median, and weighted mode methods:” |
| 13 | **Sensitivity analyses and additional analyses** |  |  |  |
|  | a) | Report any sensitivity analyses to assess the robustness of the main results to violations of the assumptions | 21-23 | Section “Interpretation of the results indicative of causality for inversed variant weighted (IVW), MR-Egger, weighted median, and weighted mode methods:” |
|  | b) | Report results from other sensitivity analyses or additional analyses | 21-23 | Section “Interpretation of the results indicative of causality for inversed variant weighted (IVW), MR-Egger, weighted median, and weighted mode methods:” |
|  | c) | Report any assessment of direction of causal relationship (e.g., bidirectional MR) | 8 | “We used the LHC-MR method to estimate bidirectional causal effects between abdominal obesity and smoking traits. Our results showed a positive causal effect of WHR on lifetime smoking (0.08 [0.04, 0.13]), WC on smoking initiation (0.12 [0.04, 0.21]) and lifetime smoking (0.14 [0.03, 0.25]), and HC on smoking initiation (0.11 [0.03, 0.2]) and lifetime smoking (0.10 [0.04, 0.16]) (Figure 4, Supplementary Table 5).” |
|  | d) | When relevant, report and compare with estimates from non-MR analyses |  | Not relevant |
|  | e) | Consider additional plots to visualize results (e.g., leave-one-out analyses) | 27 | Supplementary Figure 3 |
|  | **DISCUSSION** |  |  |  |
| 14 | **Key results** | Summarize key results with reference to study objectives | 9 | “Our study showed that smoking initiation and lifetime smoking may causally increase abdominal adipsoity as indicated by higher WHR and WHRadjBMI. Analyses for fat depot volumes indicated that visceral fat increases relatively more than abdominal subcutaneous fat. While we found no evidence of an association between smoking heaviness and abdominal fat distribution, our reverse causal analysis indicated that higher abdominal adiposity may causally increase smoking heaviness.” |
| 15 | **Limitations** | Discuss limitations of the study, taking into account the validity of the IV assumptions, other sources of potential bias, and imprecision. Discuss both direction and magnitude of any potential bias and any efforts to address them | 10 | “However, there are also some limitations to our study. Despite performing several sensitivity analyses, we cannot completely rule out the potential influence of residual pleiotropy on our causal estimates. Furthermore, the sample size for body fat distribution in current smokers was relatively small, limiting the statistical power of our smoking heaviness analysis compared to the larger sample sizes used for smoking initiation and lifetime smoking. Finally, our population was restricted to individuals of European genetic ancestry, and thus the findings may not be generalizable to other populations.” |
| 16 | **Interpretation** |  |  |  |
|  | a) | Meaning: Give a cautious overall interpretation of results in the context of their limitations and in comparison with other studies | 10 | “In conclusion, smoking initiation and lifetime smoking may causally increase abdominal and particularly visceral fat. Thus, public health efforts to prevent and reduce smoking may also help reduce abdominal fat and the risk of related chronic diseases.” |
|  | b) | Mechanism: Discuss underlying biological mechanisms that could drive a potential causal relationship between the investigated exposure and the outcome, and whether the gene-environment equivalence assumption is reasonable. Use causal language carefully, clarifying that IV estimates may provide causal effects only under certain assumptions | 10 | “Smoking may increase abdominal fat by increasing either visceral fat or abdominal subcutaneous fat. Our genetic association findings for MRI-based adipose depot volumes suggested that the effect of smoking on abdominal adiposity is mainly driven by an increase in VAT and less the increase in ASAT. These results are in agreement with previous observational studies that have shown a positive association between smoking and higher levels of VAT compared to non-smokers [6, 50]. As the amount of VAT is closely connected to the development of cardiometabolic disease, the observed changes are likely to increase cardiometabolic risk.”  “It has been suggested that smoking can affect body fat distribution through changes in cortisol and sex hormone levels [42, 52-54]. Smokers have increased levels of cortisol which is linked to insulin resistance and abdominal fat [55]. Smokers have also increased levels oestradiol, testosterone, and sex hormone-binding globulin, which may influence abdominal fat. However, our genetic association results did not support a role of either cortisol or sex hormones in the relationship between smoking and abdominal adiposity.” |
|  | c) | Clinical relevance: Discuss whether the results have clinical or public policy relevance, and to what extent they inform effect sizes of possible interventions | 10 | “As the amount of VAT is closely connected to the development of cardiometabolic disease, the observed changes are likely to increase cardiometabolic risk.”  “Thus, public health efforts to prevent and reduce smoking may also help reduce abdominal fat and the risk of related chronic diseases.” |
| 17 | **Generalizability** | Discuss the generalizability of the study results (a) to other populations, (b) across other exposure periods/timings, and (c) across other levels of exposure | 10 | “Furthermore, the sample size for body fat distribution in current smokers was relatively small, limiting the statistical power of our smoking heaviness analysis compared to the larger sample sizes used for smoking initiation and lifetime smoking. Finally, our population was restricted to individuals of European genetic ancestry, and thus the findings may not be generalizable to other populations.” |
|  | **OTHER INFORMATION** |  |  |  |
| 18 | **Funding** | Describe sources of funding and the role of funders in the present study and, if applicable, sources of funding for the databases and original study or studies on which the present study is based | 11 | “The present study was conducted independently without any involvement or influence from any of the fundings agencies. Novo Nordisk Foundation Center for Basic Metabolic Research is an independent research center at the University of Copenhagen partially funded by an unrestricted donation from the Novo Nordisk Foundation (NNF18CC0034900). Germán D. Carrasquilla was supported by a grant from the Danish Diabetes Academy that is funded by the Novo Nordisk Foundation (NNF17SA0031406), and from the European Union’s Horizon 2020 research and innovation programme under the Marie Sklodowska-Curie grant agreement No 846502. Tuomas O. Kilpeläinen was supported by the grants NNF17OC0026848, NNF21SA0072102, and NNF22OC0074128 from the Novo Nordisk Foundation.” |
| 19 | **Data and data sharing** | Provide the data used to perform all analyses or report where and how the data can be accessed, and reference these sources in the article. Provide the statistical code needed to reproduce the results in the article, or report whether the code is publicly accessible and if so, where | 6 | “The present study used public GWAS summary-level data and did not have direct contact with study participants. Therefore, no direct ethical approval was needed. The code and curated data for the current analysis are available at https://github.com/MarioGuCBMR/MR_smoking_abdominal_adiposity.” |
| 20 | **Conflicts of Interest** | All authors should declare all potential conflicts of interest | 1 | “Declaration of interests: none” Following journal regulations. |

This checklist is copyrighted by the Equator Network under the Creative Commons Attribution 3.0 Unported (CC BY 3.0) license.

1.	Skrivankova VW, Richmond RC, Woolf BAR, Yarmolinsky J, Davies NM, Swanson SA, et al. Strengthening the Reporting of Observational Studies in Epidemiology using Mendelian Randomization (STROBE-MR) Statement. JAMA. 2021;under review.

2.	Skrivankova VW, Richmond RC, Woolf BAR, Davies NM, Swanson SA, VanderWeele TJ, et al. Strengthening the Reporting of Observational Studies in Epidemiology using Mendelian Randomisation (STROBE-MR): Explanation and Elaboration. BMJ. 2021;375:n2233.
